# GENETIC ARCHITECTURE OF DIVERGENCE: THE SELFING SYNDROME IN *IPOMOEA LACUNOSA*

**DOI:** 10.1101/2021.01.14.426677

**Authors:** J. L. Rifkin, G. Cao, M. D. Rausher

## Abstract

**Premise of the study:** Highly selfing plant species frequently display a distinctive suite of traits termed the “selfing syndrome.” This study tests the hypothesis that these traits are grouped into correlated evolutionary modules and determines the degree of independence between such modules.

**Methods:** We evaluated phenotypic correlations and QTL overlaps in F2 offspring of a cross between the morning glories *Ipomoea lacunosa* and *I. cordatotriloba* and investigated how traits clustered into modules at both the phenotypic and genetic level. We then compared our findings to other QTL studies of the selfing syndrome.

**Key results:** In the *I. lacunosa* selfing syndrome, traits group into modules that display correlated evolution within but not between modules. QTL overlap predicts phenotypic correlations, and QTLs affecting the same trait module are significantly physically clustered in the genome. The genetic architecture of the selfing syndrome varies across systems, but the pattern of stronger within-than between-module correlation is widespread.

**Conclusions:** The genetic architecture we observe in the selfing syndrome is consistent with a growing understanding of floral morphological integration achieved via pleiotropy in clustered traits. This view of floral evolution is consistent with resource limitation or predation driving the evolution of the selfing syndrome, but invites further research into both the selective causes of the selfing syndrome and how genetic architecture itself evolves in response to changes in mating system.

## INTRODUCTION

The transition from outcrossing to selfing has occurred repeatedly in angiosperms (Barrett, 2002), and is often associated with additional trait changes beyond the shift in mating system. Self-pollinating species generally produce smaller flowers than outcrossing relatives, often with reduced color, nectar, pollen, and scent. They also frequently differ from outcrossing relatives in non-floral traits, including inflorescence structure and phenology. By analogy with biotic pollination syndromes, these changes are collectively termed the “selfing syndrome” (Ornduff, 1969; Sicard and Lenhard, 2011; Woźniak and Sicard, 2018).

Unlike biotic pollination syndromes, the selfing syndrome does not evolve in response to an obvious agent of selection. Nevertheless, there are reasons to believe that selection drives its evolution. Repeated convergent evolution is itself often taken as evidence of repeated responses to similar selective pressures (Stern, 2013). In addition, Qst-Fst studies (Duncan and Rausher, 2013a; Rifkin, Liao, et al., 2019) and molecular analyses (Tsuchimatsu et al., 2020) indicate that at least some selfing-syndrome traits have evolved in response to natural selection. However, for most selfing syndrome traits, no such evidence exists.

There are three possible explanations for the evolution of these traits. They may have been favored by selection acting directly on them. Alternatively, they may have evolved through a correlated response to selection on other, genetically correlated traits. Finally, genetic drift could be largely responsible for their evolution. These explanations are not necessarily independent or mutually exclusive. The degree of trait correlation also shapes the way selection can effect trait change: when traits are uncorrelated, adaptive change requires selection to act on each trait individually, but if several traits are positively correlated, selection on just one of them can cause all of them to change in the same direction. Determining the degree to which selfing syndrome traits are genetically independent thus informs us about the nature of selection required to produce the syndrome.

Correlated evolution is a plausible explanation for selfing-syndrome changes. Floral traits often evolve in a strikingly coordinated fashion, likely due to pleiotropic effects of substitutions in loci underlying selected traits (Smith, 2015; Wessinger and Hileman, 2016). Theory predicts that trait correlations can either facilitate or impede evolutionary change (Lande and Arnold, 1983), and experimental evolution studies of selfing syndrome traits suggest facilitation may be common. For example, selection for smaller flowers in *Eichornia paniculata* (Worley and Barrett, 2000) also reduces nectar production, and in *Phlox* selection on corolla size produces correlated changes in tube length and anther-stigma separation (Lendvai and Levin, 2003).

QTL studies of the selfing syndrome also tend to support this pattern of trait correlation, but reveal wide variation in genetic architecture across systems. At one extreme, QTLs affecting traits included in the selfing syndrome of *Capsella rubella* (Sicard et al., 2011; Slotte et al., 2012) are highly overlapping and localized to few regions of the genome. At the opposite extreme, the selfing syndrome of *Mimulus* is controlled by numerous loci that, although highly overlapping, are scattered widely across the genome (Lin and Ritland, 1997; Fishman et al., 2002, 2014). In *Solanum*, both the number of regions and the degree of overlap are low (Bernacchi and Tanksley, 1997; Georgiady et al., 2002). However, these previous QTL studies have generally focused primarily on floral morphology and reproductive traits, and have not consistently included the nectar, pollen, or phenological traits also typically associated with the selfing syndrome. Consequently, the extent to which these traits are genetically independent of floral morphological traits, and thus evolve independently, remains an open question.

In this study, we examine these issues in the highly selfing *Ipomoea lacunosa* and its sister species, *I. cordatotriloba*. Specifically, we examine the extent to which a broad array of selfing-syndrome traits, including floral morphology, inflorescence structure, pollen traits, nectar production, and phenology, have evolved in either an independent or a correlated manner in *I. lacunosa*. We apply two approaches. First, we assess trait phenotypic correlations in an F2 population resulting from a cross between two closely related species, one of which exhibits the selfing syndrome. Such correlations are expected to reflect underlying levels of pleiotropy in loci involved in selfing syndrome evolution. We identify evolutionary modules from these correlations. Second, we determine whether the co-localization of quantitative trait loci (QTLs) for selfing-syndrome traits is consistent with pleiotropy. Specifically, we ask whether QTL overlap is higher within modules than between modules, as would be expected for independent evolutionary modules (Wagner and Altenberg, 1996; Brandon, 1999; Armbruster et al., 2014). We then consider whether results of the two approaches provide a consistent picture of the genetic architecture of divergence for the selfing syndrome.

The genetic architecture of divergence also offers insight into which selective forces likely shaped the selfing syndrome of a given species. Different selective explanations for the evolution of the selfing assume different amounts of correlation between traits (Sicard and Lenhard, 2011), with some requiring high correlation and others allowing independent trait evolution. We apply our findings about the degree of overlap in *I. lacunosa* to infer which possible selective causes were likely at play in the divergence of these species. Finally, we place our results in the context of selfing syndrome evolution in angiosperms more generally by performing a quantitative review of QTL overlap in studies of the selfing syndrome. Specifically, we ask how genetic architecture, reflected in patterns of overlap, varies among taxa. These general trends offer the opportunity for inferences about which selective factors most frequently drive the evolution of the selfing syndrome.

## METHODS

### Study system

*Ipomoea lacunosa* and *I. cordatotriloba* are weeds in series *Batatas* of the Convolvulaceae. They occur in overlapping ranges in the southeastern United States (USDA and NRCS, 2017). *Ipomoea lacunosa* extends north and east as far as Canada, and *I. cordatotriloba* extends south and west to Mexico. *Ipomoea lacunosa* is highly selfing (>0.95), while *I. cordatotriloba* has a mixed mating system with highly selfing, highly outcrossing, and intermediate populations (Duncan and Rausher, 2013b). Although vegetatively similar, the two species are genetically distinct (Rifkin, Castillo, et al., 2019) and have dramatically different flowers: *I. cordatotriloba* produces medium sized (corolla length 32.1 ± 4.1mm) purple flowers, while *I. lacunosa* produces small (corolla length 20.0 ± 1.4 mm) white flowers (Rifkin, Liao, et al., 2019). Consistent with the selfing syndrome, *I. lacunosa* also produces less nectar and pollen and exhibits reduced inflorescences and faster early growth (Rifkin, Liao, et al., 2019). Differences in corolla size and in nectar production are due to selection (Duncan and Rausher, 2013a; Rifkin, Liao, et al., 2019). Both species are diploid, with small genomes (*I. lacunosa* 497mb, *I. cordatotriloba* 525 mb (Duncan and Rausher, 2013b)) and 15 pairs of chromosomes (Nakajima, 1963).

### Generation of mapping population

We generated a mapping population from seeds collected from a wild *I. lacunosa* population in Kinston, North Carolina (lat. 35.23971, long. -77.57392) and a wild *I. cordatotriloba* population in Conway, South Carolina (lat. 33.94713, long. -79.01940) (Duncan and Rausher, 2013a). Wild individuals were inbred for four generations to develop our lab strains LPR (“Lacunosa ProTruck”) and CAA (“Cordatotriloba Double-A Lane”). The *I. cordatotriloba* population is sympatric with *I. lacunosa* and shows evidence of substantial introgression from that species (Rifkin, Castillo, et al., 2019), although this was unknown at the time these individuals were chosen. Despite this introgression, traits differ substantially between individuals of the two species used in our crosses (Table 1; (Rifkin, Castillo, et al., 2019; Rifkin, Liao, et al., 2019)). Crosses were performed between clones of these individuals by emasculating *I. cordatotriloba* buds and pollinating with whole *I. lacunosa* flowers. We generated seven F1 individuals from this cross, cloned them via cuttings, and allowed them to self to generate F2 seeds. 508 F2 seeds from a single F1 individual (“CL5”) formed our mapping population.

**Table 1.** Mean trait values from grandparent genotype clones (floral, color, inflorescence, pollen, and nectar traits) or selfed offspring of grandparents (early growth life-history traits) in 2014 growout (2013 values in parentheses). Module is trait group identified by cluster analysis. Asterisks indicate that F2 mean differs significantly from midparent value, or that F1 differed from midparent value in 2013 growout, following a false discovery rate correction.

| <u>Anatomical</u> | <u>Cluster</u> | <u>Trait (n F2s</u> |  | <u>*I. lacunosa*</u> | <u>*I. cordatotriloba*</u> |  |  |
| --- | --- | --- | --- | --- | --- | --- | --- |
| <u>Module</u> | <u>Module</u> | <u>phenotyped)</u> | <u>Acronym</u> | <u>grandparent</u> | <u>grandparent</u> | <u>F1 parent</u> | <u>F2 mean</u> |
| Color | -- | Color (356) | COL | 0.00 | 1.00 | 1.00 | 0.79* |
| Floral |  |  |  | 18.78 | 30.74 | 24.55 |  |
| morphology | 1 | Corolla Length (356) | CL | (16.94) | (28.37) | (22.84) | 26.65* |
| Floral |  | Corolla Shape 1 (CL / |  |  |  |  |  |
| morphology | 1 | CW) (356) | LW | 1.35 | 1.19 | 1.20 | 1.26* |
| Floral |  | Corolla Shape 2 (TL/ |  |  |  |  |  |
| morphology | 1 | CL) (356) | TLL | 1.13 | 1.16 | 1.15 | 1.15* |
| Floral |  | Corolla Tissue Length |  |  |  |  |  |
| morphology | 1 | (356) | TL | 21.15 | 35.53 | 28.22 | 30.73* |
| Floral |  |  |  | 13.97 | 26.01 | 20.47 |  |
| morphology | 1 | Corolla Width (356) | CW | (13.56) | (24.64) | (19.72) | 21.32* |
| Floral |  |  |  | 11.16 | 17.52 | 15.58* | 15.91* |
| morphology | 1 | Style Length (356) | SL | (9.67) | (14.64) | (13.39)* |  |
| Inflorescence | 2 | Cyme Length (356) | CY | 15.88 | 20.92 | 16.05 | 23.02* |
| Inflorescence |  | Flowers and Buds on |  |  |  |  |  |
|  | 2 | Cyme (356) | FC | 1.15 | 1.41 | 1.17 | 1.42* |
| Phenology |  |  |  | 470.61 | 297.28 |  | 442.08* |
|  | 5 | Height (424) | H | (492.91) | (334.90) | (561.92) |  |
| Phenology |  | Length of First |  | 15.20 | 6.47 |  |  |
|  | 3 | Internode (424) | INT1 | (13.03) | (6.33) | (10.15) | 11.56* |
| Phenology |  | Length of Second |  | 17.02 | 7.72 |  |  |
|  | 3 | Internode (424) | INT2 | (22.36) | (6.86) | (17.85) | 16.38* |
| Phenology |  | Length of Third |  | 67.41 | 22.72 |  |  |
|  | 3 | Internode (424) | INT3 | (79.48) | (26.40) | (78.31)* | 63.27* |
| Phenology |  | Number of Leaves |  | 5.74 | 6.19 |  |  |
|  | 5 | (424) | NL | (7.36) | (8.59) | (9.92) | 5.96 |
| Phenology | 2 | Flowering Date (356) | FD | N/A | N/A | N/A | 158.62 |
| Phenology |  | Number of Flowers |  |  |  |  |  |
|  |  | Measured (419) | NF | N/A | N/A | N/A | 3.14 |
| Phenology |  | Number of Flowers |  |  |  |  |  |
|  | 4<sup>1</sup> | Open (356) | FPD | N/A | N/A | N/A | 1.29 |
| Phenology |  | Ever Flowered (419) | EF | N/A | N/A | N/A | 0.834 |
| Phenology |  | First Leaf Opening |  | 11.53 | 11.22 |  |  |
|  | 5 | (419) | L1 | (13.54) | (12.86) | (12.00)* | 11.62 |
| Phenology | 5 | Second Leaf Opening | L2 | 13.78 | 13.07 |  | 13.60 |
|  |  | (416) |  | (15.95) | (15.09) | (14.31) |  |
| Phenology |  | Third Leaf Opening |  | 16.04 | 15.50 |  | 15.88 |
|  | 5 | (401) | L3 | (18.64) | (18.10) | (17.00) |  |
| Nectar | 1(2)<sup>1</sup> | Nectar Volume (356) | NVL | 0.00 | 1.31 | 0.40 | 0.21* |
| Pollen | 4 | Pollen Count 1, (69) | P1 | 140.00 | 706.00 | 323.50 | 204.48* |
| Pollen | 4 | Pollen Count 2, (110) | P2 | 450.00 | 946.25 | 251.25 | 350.03* |
| Pollen | 4 | Pollen Size, (69) | PS | 69.54 | 81.29 | 71.85 | 69.32* |
|  | 4<sup>1</sup> | Herkogamy<sup>2</sup> (356) | HK | -0.08 | -0.03 | 0.00 | 0.01* |

<sup>1</sup> Placed in different modules by different cluster analyses. Module not in parentheses is the module the trait clustered with in the majority of clustering analyses.
<sup>2</sup> Removed from downstream analyses because of lack of segregating variation in F2s

Plants were germinated in the spring of 2014 and flowered from July 2014 through early spring of 2015. We scarified seeds and germinated them on filter paper in petri dishes in the dark. Seeds that had not swelled within two days were re-scarified. Germinated seeds were transferred to individual 4” plastic pots filled with Fafard 4P soil and grown in 16-hour days in a growth room for six weeks at 80°F. The light cycle was changed to 13 hours of light and a temperature of 65°F for two weeks to induce flowering, after which plants were transferred to the Duke Research Greenhouse and maintained there under greenhouse conditions.

### Phenotyping and trait modules

#### Phenology traits

The selfing syndrome may include phenological and life-history as well as floral traits (Snell and Aarssen, 2005), and *I. lacunosa* exhibits faster early growth than *I. cordatotriloba* (Rifkin, Liao, et al., 2019). We measured eight early growth traits (days to opening of first three leaves: L1, L2, L3; total height on day 21 with the vine unwound to its full length: H; number of leaves on day 21: NL; length of first three internodes on day 21: INT1, INT2, INT3) and three floral phenology traits (average floral measurement date: FD; whether any flowers were produced: EF; total number of flowers measured: NF) in 424 F2 hybrids (see Table 1). EF and NF were included in analyses because we observed considerable variation in both traits: during the study period, some plants produced no flowers, while others flowered continuously. Although we generally stopped measuring after 5 flowers had been sampled, we believe these measurements offer a proxy for genetic variation in the mapping population reflecting divergence in phenology between the two species. We measured early growth traits in selfed offspring of the grandparents of the mapping population in the same growout as the F2 hybrids (N=46 *I. lacunosa*, 47 *I. cordatotriloba*) but did not measure floral phenotypes in these plants because of space constraints. These traits were also measured in F1 hybrids and in the selfed offspring of the grandparents of the mapping population in a separate growout in the summer of 2013 (N=58 *I. lacunosa*, 58 *I. cordatotriloba*, 13 F1). Measurements were consistent between these growouts, with the exception of number of leaves, which was measured on day 23 rather than day 21 in the 2013 growout.

We measured total height and internode length with a rule and scored leaf opening as the time when the focal leaf unfolded to an angle of greater than 90°. Floral phenology (FD) was scored as average flower measurement date rather than days to first flower because the fast-growing vines required frequent pruning and the first flower measured was not necessarily the first flower produced but plants were measured soon after they began flowering. Because only a subset of the F2 individuals ever flowered, we scored whether a plant ever produced flowers as a binary trait. Of the 507 seeds that germinated, 424 survived transplant and were included in downstream analyses.

#### Floral and inflorescence traits

We measured flowers between 9:00 and 12:00 every morning. Only one flower was measured per individual per day. Because flowers last for only a single day, flower age is unlikely to affect our measurements. Floral traits were measured simultaneously in clones of each grandparent and the F1 parent of the mapping population. We thus had simultaneous floral measurements from only a single genotype for grandparent and F1 floral traits. 357 F2 individuals flowered and were phenotyped for ten floral and inflorescence traits (color: COL; total number of flowers open on plant containing measured flower: FD; length of cyme containing measured flower: CY; number of flowers or buds on cyme containing measured flower: FC; corolla length: CL; corolla width: CW; corolla tissue length: CTL; herkogamy: HK; nectar volume: NVL; and style length:SL; see Appendix S1 for measurement details). We also quantified pollen number (P1, P2) and size (PS) in a subset of F2 individuals. See supplementary methods for measurement details. Per-individual trait means were calculated in R using **dplyr** functions (Wickham et al., 2020) and pairwise trait correlations were calculated using the package HMisc (Harrell Jr et al., 2016) with a Bonferroni correction for multiple comparisons.

### Identifying trait modules

To assemble a selfing syndrome, traits that encompass a range of tissue types and functions evolve. To investigate their joint evolution, we grouped traits into modules and determined the amount of correlation within and between these modules. We took two approaches towards grouping our traits into modules. First, we identified modules using cluster analysis. Second, for comparison with previous studies and with the clustering-based analyses, we grouped them into anatomical modules based on organ or tissue type. The anatomically based modules we used were floral morphology, inflorescence, nectar, pollen, and phenology. We chose these groupings because all of them, with the exception of nectar, included multiple traits that are likely to be under selection in the same context, and some of them have been previously shown to also reflect genetically correlated groupings in other systems (Smith, 2015).

Our cluster analysis relied on the definition of a trait module as a group of traits that are highly genetically correlated with each other but have low genetic correlations with traits in other modules (Wagner and Altenberg, 1996; Brandon, 1999; Armbruster et al., 2014). In the context of trait divergence between species, a trait module is a group of traits that are highly genetically correlated in an F2 mapping population but show little correlation with traits in other modules. A module thus reflects divergent genes with pleiotropic effects that affect primarily traits in a single module. To the extent that phenotypic correlations in a mapping population reflect underlying genetic correlations, modules can be identified using cluster analysis on phenotypic correlations (Cheverud, 1988; Roff, 1995; Waitt and Levin, 1998; Feng et al., 2019).

For our cluster analysis, we used all measured traits in our F2 mapping population (Table 1) except color. The correlation matrix was first converted to a distance matrix by the transformation *D*_*ij*_ = 1 – *r*_*ij*_^2^, where *D*_*ij*_ is the distance between traits *i* and *j*, and *r*_*ij*_ is the phenotypic correlation between those traits. We then performed a cluster analysis on the distance matrix using three different algorithms implemented by the SAS statistical software package version 9.4 (SAS Instititute, 2013): the Averaging, Ward, and McQuitty algorithms. Modules were identified as corresponding to the five lowest-level trait groupings in the cluster analyses.

If modules have been correctly identified, we expect average correlations between traits within modules to be larger than the average correlations between traits in different modules. To evaluate this expectation, we calculated the average within- and between-module correlation (and their standard errors) for each module. We also used a permutation test (1,000 replicate permutations), in which we permuted which correlations were assigned to which trait pairs, to evaluate whether the average within-module correlation (averaged over all modules) was greater than the average between-module correlation (averaged over all module pairs). These analyses were performed by an APL program written by MDR (all APL programs available at https://github.com/joannarifkin/Ipomoea_QTL/tree/main/APL_scripts).

### Linkage map construction

We generated markers from our mapping population using double-digest restriction-site associated DNA sequencing (ddRADseq; Peterson et al., 2012) and sequenced them using the Illumina HiSeq platform (see Appendix S2 for details of extraction, restriction enzyme digest, barcoding, amplification, and sequencing). After controlling for contamination, we ultimately included 396 F2 individuals in our linkage map and subsequent QTL analysis. A linkage map was constructed using Lep-Map3 (Rastas et al., 2013) from reads aligned to the *I. lacunosa* draft assembly (see Appendix S3 for details). We then used Lep-Anchor (Rastas, 2020) to relate our linkage map to our draft assembly and generate marker orders that incorporated the physical and linkage estimates, and a custom Python script to convert marker names to positions on chromosomes. The final map included 6124 markers in 15 linkage groups. Scripts are available on Github (https://github.com/joannarifkin/Ipomoea_QTL).

### QTL mapping

We generated genotype data for QTL mapping using Lep-Map3’s map2genotypes function and used custom scripts to identify grandparent genotypes from the raw Lep-Map3 genotypes. Single-QTL analyses were performed in QTL2 (Broman et al., 2019). Traits that were normally distributed or binary were kept unchanged, while traits with highly skewed distributions were transformed and in some cases split; see Appendix S4 for details. After splitting and transformation, we identified QTLs for a total of 27 phenotypes in 396 phenotyped and genotyped individuals (Table 1, Appendix S5). To identify QTLs, we used the scan1 function in QTL2 with either the default or binary model (see Appendix S5 for details). We determined significance cutoffs using permutation analysis (1000 replicates) at both genome-wide and chromosome-wide levels. Confidence intervals were estimated as 1.5-LOD intervals. Additive and dominance effects were estimated using the scan1coef function in QTL2. QTLs were categorized as over- or under-dominant if the heterozygote value was outside of the range of the homozygote values. Scripts are available on Github (https://github.com/joannarifkin/Ipomoea_QTL).

### QTL overlap

To quantitatively describe the genetic architecture underlying the selfing syndrome, we examined the degree to which QTLs for different traits co-localized, and thus are potentially represented by the same underlying genetic variant. A pair of QTLs was considered to co-localize if they had overlapping 1.5 LOD confidence intervals. As an indicator of average overlap for a pair of traits, we used the Jaccard Index: J = I/(A + B – I), where I = number of co-localizing QTLs, A is the total number of QTLs for the first trait, and B is the total number of QTLs for the second trait. Average within- and between-module QTL overlap (and standard errors) were calculated in an APL program written by MDR. To determine whether within-module average overlaps and between-module average overlaps differed significantly, we performed a permutation test (1,000 replicates) in which the QTLs were randomized with respect to traits. We performed these analyses separately for QTLs that were significant only genome-wide and QTLs that were significant both genome- and chromosome-wide.

We also performed an analysis to determine whether QTLs within modules are significantly concentrated spatially compared to random placement of QTLs, as would be expected if QTL overlap indicates pleiotropy. For traits within a module, we randomized the position of QTLs by weighting 50-kb bins along chromosomes by the number of genes within those bins then choosing randomly among the weighted bins. We identified genes using a newly generated annotation (Appendix S6). The position of the QTL within the selected bin was chosen randomly. The size of each repositioned QTL was maintained, only its position was changed. Once all QTLs associated with a module were positioned, we calculated the average QTL overlap across module trait pairs as described above. This randomization process was repeated 1,000 times to determine the proportion of average QTL overlap values greater than the observed value. In addition to performing this analysis on each module separately, we also performed an analogous analysis for the overall average QTL overlap for all within-module trait pairs. These analyses were performed using APL scripts written by MDR (available on Github: https://github.com/joannarifkin/Ipomoea_QTL).

To determine whether patterns identified in our analysis of trait correlations were consistent with those identified by our QTL analysis, we performed two types of analysis, each using either all QTLs or only GWS QTLs. The first analysis asked whether phenotypic correlations between trait pairs were correlated with Jaccard indices of overlap. Significance was evaluated by permuting the phenotypic correlation 1,000 times. In the second analysis, we asked whether phenotypic correlations were correlated with predicted genetic correlations, where genetic correlations were predicted from QTL properties following a modified version of the approach of Gardner and Latta (2007). Significance was evaluated by permuting QTLs among traits (see Appendix S7).

### QTL overlap review

To compare our results to previous studies that examined the genetic architecture of the selfing syndrome, we identified appropriate studies from reviews of genetic architecture of floral changes and from the search terms “selfing syndrome” and “QTL” on Google Scholar, Web of Science, and ProQuest. For each study, we extracted all QTLs as a table and calculated the Jaccard Index of overlap for all trait pairs. We then categorized traits into the following trait modules: floral morphology, nectar, inflorescence structure, phenology, pollen, reproductive, or vegetative. Reproductive traits, which were not included in our QTL study, consisted of seed number, fruit, and compatibility traits. We also treated herkogamy as a reproductive trait. We selected these categories because QTL studies of floral trait evolution generally suggest that these groupings reflect biological realities and to facilitate comparison with our anatomical modules (Ashman and Majetic, 2006; Smith, 2015; Feng et al., 2019).

## RESULTS

### Phenotypic distributions

The parents of our mapping population reflected known differences in selfing-syndrome traits between *Ipomoea lacunosa* and *I. cordatotriloba* ((Rifkin, Liao, et al., 2019), Table 1). However, both parents exhibited similarly low herkogamy, perhaps because the *I. cordatotriloba* parent of the mapping population was collected from a population sympatric with *I. lacunosa*; we therefore did not include herkogamy in subsequent analyses. In the F1 and F2 offspring, we observed a general tendency towards *I. cordatotriloba* dominance in floral traits and *I. lacunosa* dominance in phenological traits (Table 1). Finally, color is known to be under single-locus control in *I. lacunosa* (Duncan and Rausher, 2020) and, consistent with this, F2s produced purple and white flowers in a ratio that did not differ from 3:1 (χ^2^ = 3.8352, P = 0.0501).

### Modules identified by cluster analysis

There was substantial variation in the magnitude of the F2 phenotypic correlations, which ranged from 0 to 0.965. Half of the correlations (115) were negative, while the other half (116) were positive (Appendix S4, Appendix S8, Appendix S9). Based on these correlations, the three cluster algorithms used generally produced a consistent set of modules (Table 1; Appendix S10), which were broadly similar to the anatomical modules of floral morphology, phenology, inflorescence traits, nectar, and pollen. This suggests that in the absence of more complete genetic or phenotypic data, anatomical groupings reflect underlying genetic groupings. Module 1 consists of all floral dimension traits and nectar volume. Module 2 includes both inflorescence traits and the phenological trait of flowering date. Module 3 consists of a subset of phenological traits (all internode lengths). Pollen traits cluster in module 4, along with the phenological trait flowers per day. Finally, module 5 consists of the remaining phenological traits (dates of leaf emergence, number of leaves on day 21, height on day 21).

Three traits clustered in different modules across the three clustering algorithms: nectar volume clustered in module 1 in two analyses and in module 2 in one; flowers per day clustered with module 4 in two analyses and alone in one analysis; and herkogamy clustered with module 4 in one analysis and alone in two analyses. In subsequent analyses, we grouped traits in whichever module most frequently included them.

### Correlations within and between modules

Average trait correlations within a module are substantially higher than average trait correlations between modules (Table 2). Within-module correlations averaged 0.509, almost five times higher than between-module correlations (0.105; P<0.001, permutation test).

**Table 2.** Average pairwise trait correlations (and standard error) within (main diagonal) and between (off-diagonal) modules. Within module averages calculated as the average of correlations for all possible trait pairs within the module. Between module averages calculated as average of correlations between a trait in one module and a trait in the second module. Mean within-module correlation = 0.509. Mean between-module correlation = 0.105. Difference significant by permutation test (P < 0.001).

|  | Module 1 | Module 2 | Module 3 | Module 4 | Module 5 |
| --- | --- | --- | --- | --- | --- |
| Module 1 | 0.404 (0.057)* | 0.155 (0.027) | 0.039 (0.006) | 0.069 (0.009) | 0.055 (0.005) |
| Module 2 |  | 0.366 (0.096) | 0.050 (0.019) | 0.108 (0.026) | 0.090 (0.028) |
| Module 3 |  |  | 0.607 (0.10) | 0.126 (0.028) | 0.175 (0.041) |
| Module 4 |  |  |  | 0.342 (0.082) | 0.230 (0.030) |
| Module 5 |  |  |  |  | 0.844 (0.026) |

Nectar volume has a lower average correlation with other traits in module 1 (0.314, standard error = 0.041) than the average correlation among those traits (0.440, s.e. = 0.076), suggesting that nectar volume may not actually belong to module 1. However, this trait exhibits high correlations with corolla tissue length and corolla width (0.451 and 0.423, respectively), which are approximately equal to the average within-module correlation for module 1. Moreover, the average correlation of nectar volume with other module 1 traits is approximately equal to that of LW and TLL with other module 1 traits (0.347 and 0.313, respectively). On balance, therefore, we believe that the evidenced for removing nectar volume from module 1 is weak. Within-module correlations were strongest for modules 2 and 3. One measure of pollen number was positively correlated with pollen size; this suggests that rather than experiencing a size-number tradeoff, *I. cordatotriloba* invests more in pollen in multiple ways, as has also been shown for nectar in this species (Rifkin, Liao, et al., 2019).

With the exception of flower measurement date, which was negatively correlated with inflorescence, flower size, and nectar traits, we found almost no significant correlations between phenology and floral traits.

### Linkage map

We initially placed 6530 markers on 19 linkage groups, including fifteen large groups taken to correspond to the 15 chromosomes in the *I. lacunosa* karyotype (Nakajima, 1963). We used this map to assign the positions of 49 of the 50 scaffolds in our draft genome assembly to 15 linkage groups. We then reordered the markers according the physical order, removed markers that caused large gaps, and joined two small linkage groups to large linkage groups based on draft assembly physical order. Our final map consisted of 6124 markers on 15 LGs of between 174 and 627 markers each, with a total map length of 2222.8cm and LGs averaging 148.19cm.

Overall, the linkage map is consistent with the known karyotype of *I. lacunosa*: we identified 15 linkage groups, consistent with the 15 chromosomes. We also tested for segregation distortion in the completed map. 581 markers (9%) displayed significant segregation distortion following a Bonferroni correction. Distorted markers were concentrated on LGs 3 (117 distorted markers), 10 (126 distorted markers), 11 (176 distorted markers), and 15 (116 distorted markers).

### QTL mapping and effect sizes

We identified 50 QTLs that were significant genome-wide (GWS), and an additional 101 that were significant only at the chromosome level (CWS; Fig. 1A, B, Appendix S4). QTLs were widely scattered across the linkage groups, with 4-12 CWS and 1-8 GWS QTLs per chromosome.

**Figure 1:**
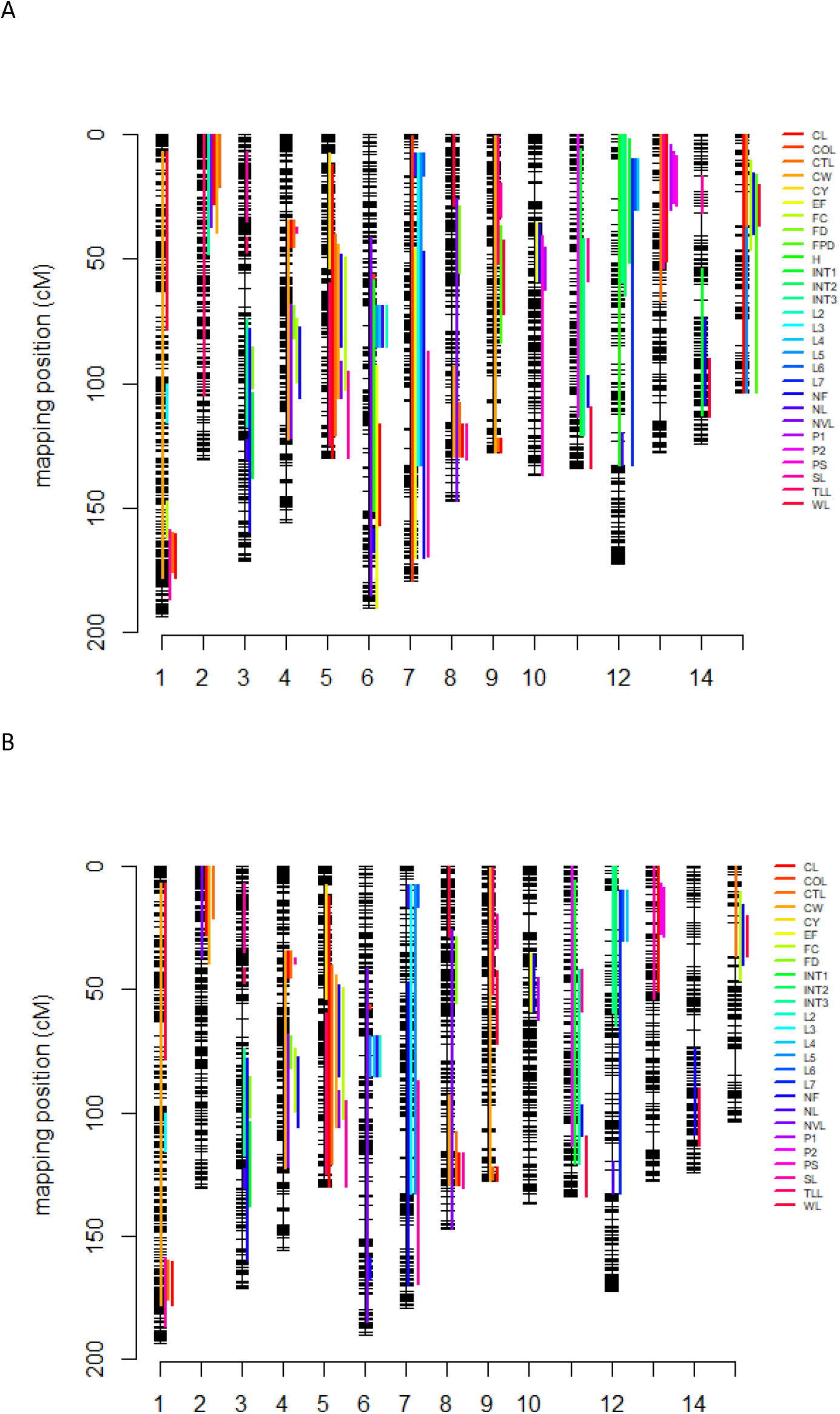
Locations of QTLs in the genome. A: All QTLs. B: GWS QTLs.

Because we believe the CWS QTLs we identified are likely real (see Discussion), we examined patterns of Percent of Variation explained (PVE) and Relative Homozygous Effect sizes (RHE) for all detected QTLs. Both total PVE and total RHE varied substantially across traits (Appendix S11).

Modules differed in average effect sizes of the QTLs associated with them (Appendix S4, Appendix S12). In general, floral morphology and nectar volume, which clustered together as module 1, had QTLs of relatively small effect (floral morphology mean RHE 0.093, maximum RHE 0.288; nectar volume mean RHE 0.061, maximum RHE 0.082). Between-species differences in these traits thus reflect the accumulation of numerous small-effect QTLs. By contrast, between-species differences in the life-history, pollen, and inflorescence traits that clustered in traits of modules 2, 4, and 5 (internode lengths, pollen traits, and inflorescence size, respectively) were on average caused by QTLs with moderate effect sizes (mean and maximum RHE > 0.1 and < 0.5), while the subset of phenology traits that clustered in module 3 were caused by substitutions of alleles with large effect (mean and maximum RHE > 0.5). In fact, many of these QTLs had RHE > 1.0, which means individually they caused a change in the trait larger than the final difference between species. The sum of the signed RHE for a trait (TRHE, Appendix S11) indicates how well the detected QTLs explain the species difference in that trait. Excluding L1 and L2, which had strongly negative TRHE values, the average proportion of parental difference explained was 0.42. Thus while a substantial number of QTL were identified, there must be undetected QTLs, presumably of smaller effect, that account for a little more than half of the observed differences between the two parents.

For both GWS QTLs and CWS QTLs, the number of QTLs that had effects in the same direction as the species difference (consistent-directional QTLs) was substantially larger than the number having effects in the opposite direction as the species difference (contra-directional QTLs; Appendix S4, Appendix S13), consistent with selection operating to fix many of these substitutions. The proportion of QTLs with consistent-directional effects was not statistically different for the two sets of QTLs (Appendix S13). Module 1 traits in particular (excluding style length) show high proportions of consistent-directional QTLs (0.95, 0.80 and 0.89 for GWS, CWS and All QTLs, respectively; Appendix S13). By contrast, for style length, the corresponding proportions are only 0.5, 0.25 and 0.4. Traits in other modules exhibit proportions that are considerably lower (0.5, 0.79, and 0.66) than for module 1 traits excluding style length.

### QTL overlap and predicted genetic correlations

QTL overlap is substantially higher within modules than between modules. Mean overlap for all traits was 0.166 for all QTL and 0.114 for GWS QTLs with genome-wide significance. For only QTLs with genome-wide significance, within-module average overlap (0.320) was significantly higher than between-module average overlap (0.037; P < 0.001, permutation test; Table 3A). An analysis using all QTLs showed similar results (Table 3B; within- and between-module average overlaps 0.320 and 0.056, respectively; P < 0.001). These analyses show that within-module overlap was at least 6 times greater than between-module overlap, indicating almost complete genetic independence of the modules.

**Table 3.** Average QTL overlap (and standard error) within and between anatomical and cluster-based modules. A. Only QTLs exhibiting genome-wide significance. Average overlap (and standard error) within modules = 0.320 (0.067). Average overlap between modules = 0.0368 (0.014). Difference significant at P < 0.001 (permutation test). B. All QTLs. Average overlap (and standard error) within modules = 0.320 (0.036). Average overlap between modules = 0.056 (0.008). Difference significant at P < 0.001 (permutation test).

**A. GWS QTLs**
|  | Module 1 | Module 2 | Module 3 | Module 4 | Module 5 |
| --- | --- | --- | --- | --- | --- |
| Module 1 | 0.201 (0.064) | 0.024 (0.024) | 0.000 (0.000) | 0.000 (0.000) | 0.000 (0.000) |
| Module 2 |  | 0.000 (0.000) | 0.000 (0.000) | 0.000 (NA) | 0.000 (0.000) |
| Module 3 |  |  | 1.000 (1.000) | 0.000 (0.000) | 0.333 (0.077) |
| Module 4 |  |  |  | 0.000 (0.000) | 0.000 (0.000) |
| Module 5 |  |  |  |  | 0.625 (0.100) |

**B. All QTLs**
|  | Module 1 | Module 2 | Module 3 | Module 4 | Module 5 |
| --- | --- | --- | --- | --- | --- |
| Module 1 | 0.299 (0.053) | 0.112 (0.026) | 0.044 (0.015) | 0.064 (0.017) | 0.025 (0.008) |
| Module 2 |  | 0.178 (0.097) | 0.019 (0.019) | 0.021 (0.021) | 0.011 (0.011) |
| Module 3 |  |  | 0.367 (0.117) | 0.033 (0.022) | 0.225 (0.060) |
| Module 4 |  |  |  | 0.222 (0.102) | 0.010 (0.010) |
| Module 5 |  |  |  |  | 0.452 (0.065) |

Randomly reshuffling QTL spatial positions throughout the genome suggests that QTLs within a module are significantly clustered spatially within the genome, as has been observed in other systems (Salih and Adelson, 2009; Kostyun et al., 2019). Using all QTLs, for each of the five modules, as well as for all modules pooled, the probability of obtaining the observed average within-module QTL overlap by random placement of QTLs in the genome was less than 0.001 (Table S5). With only GWS QTLs, modules 1 and 5, as well as the pooled average were also highly significant (P < 0.001), while module 3 was not significant (P = 0.066) and modules 2 and 4 could not be evaluated because they had QTLs for only 1 trait.

Both this analysis and that of the F2 character correlations produce similar patterns of high within-module integration and low between-module integration. We quantified this similarity in two ways. First, we calculated the correlation between pairwise trait correlations and pairwise QTL overlap. For all QTLs, this correlation was moderately high (0.64; P < 0.001), whereas for only GWS QTLs, there was little correlation (0.046) (Fig. 2A, B). Second, we examined the correlation of average trait correlations with QTL overlap over all module pairs (including a module paired with itself). Using all QTLs, this correlation is high (0.717) and significant (P = 0.005, permutation test), whereas with only GWS QTLs, the correlation is low and non-significant by a permutation test (Fig. 2C). This concordance of the two measures of integration suggests that both the F2 character correlations and the QTL overlap patterns reflect the same underlying genetic architecture of distinct modules, at least when all QTLs are considered.

**Figure 2:**
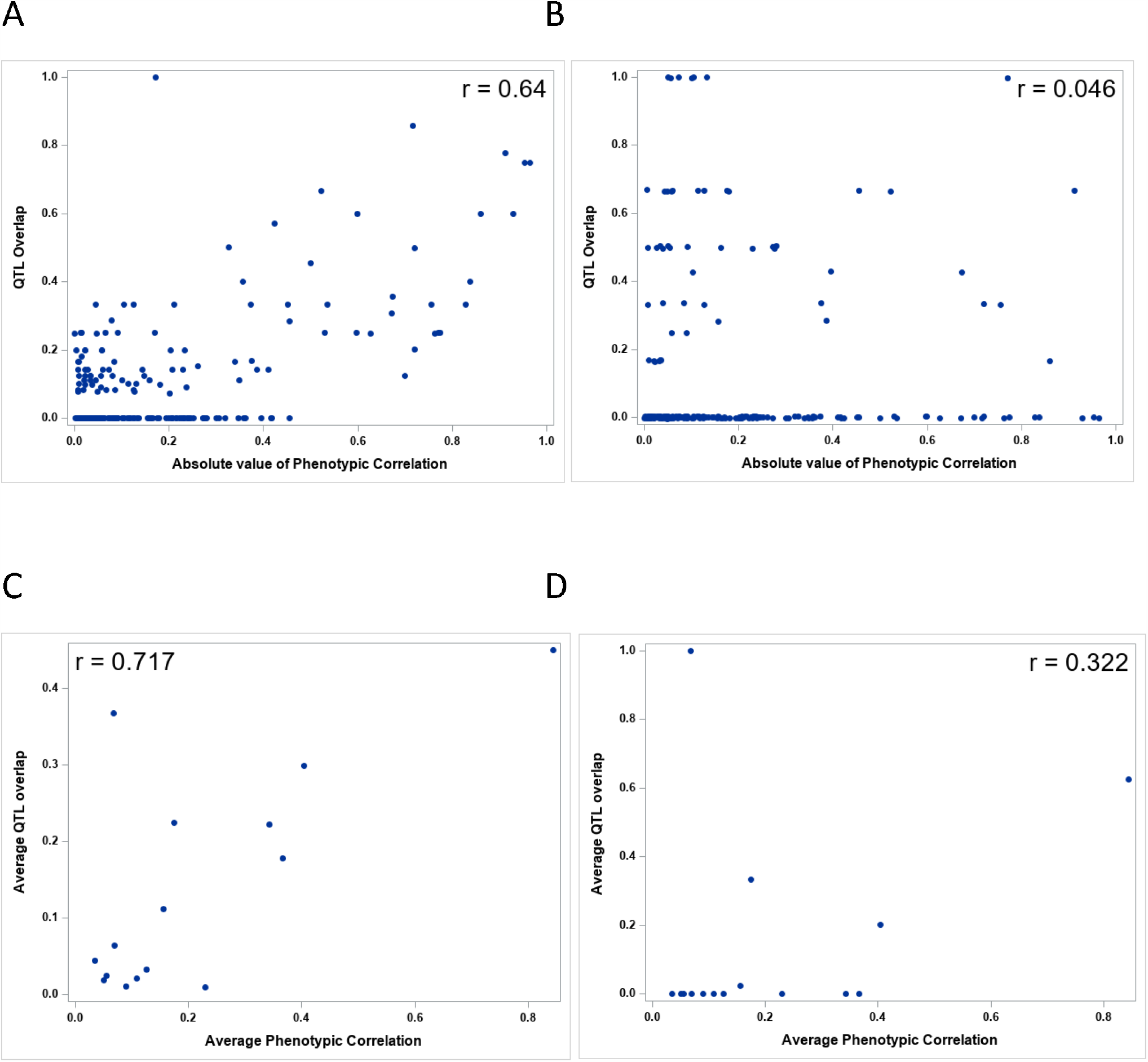
Relationship between QTL overlap and phenotypic correlations. A and B: All pairwise trait values of QTL overlap and phenotypic correlation. A. All QTLs. Correlation significant by permutation test (P < 0.001). B. Only GWS QTLs. Correlation not significant by permutation test (P = 0.233). C and D: Average within- and between-module values. C. All QTLs. Correlation significant by permutation test (P = 0.005). D. Only GWS QTLs. Correlation not significant by permutation test (P = 0.113).

Comparing the predicted genetic correlations with the observed phenotypic correlations suggests that our observed QTL overlaps captured the genetic architecture moderately effectively, as has been observed in QTL studies of other systems (Gardner and Latta, 2007). For all QTLs, the correlation between the predicted genetic correlation and the observed phenotypic correlation was 0.516, and for GWS QTLs only it was 0.546 (Fig. 3). Removing trait pairs predicted to have no genetic correlation (i.e., trait pairs with no overlapping QTLs) increased the strength of the correlations slightly to 0.554 and 0.582. All correlations are highly significant (P < 0.001, Permutation test). Using pairwise module averages produced a similar result, with a moderate correlation using all QTLs (r = 0.621, P = 0.011) and a slightly smaller correlation using only GWS QTLs (r = 0.437, P = 0.066; Fig. 3E, F).

**Figure 3:**
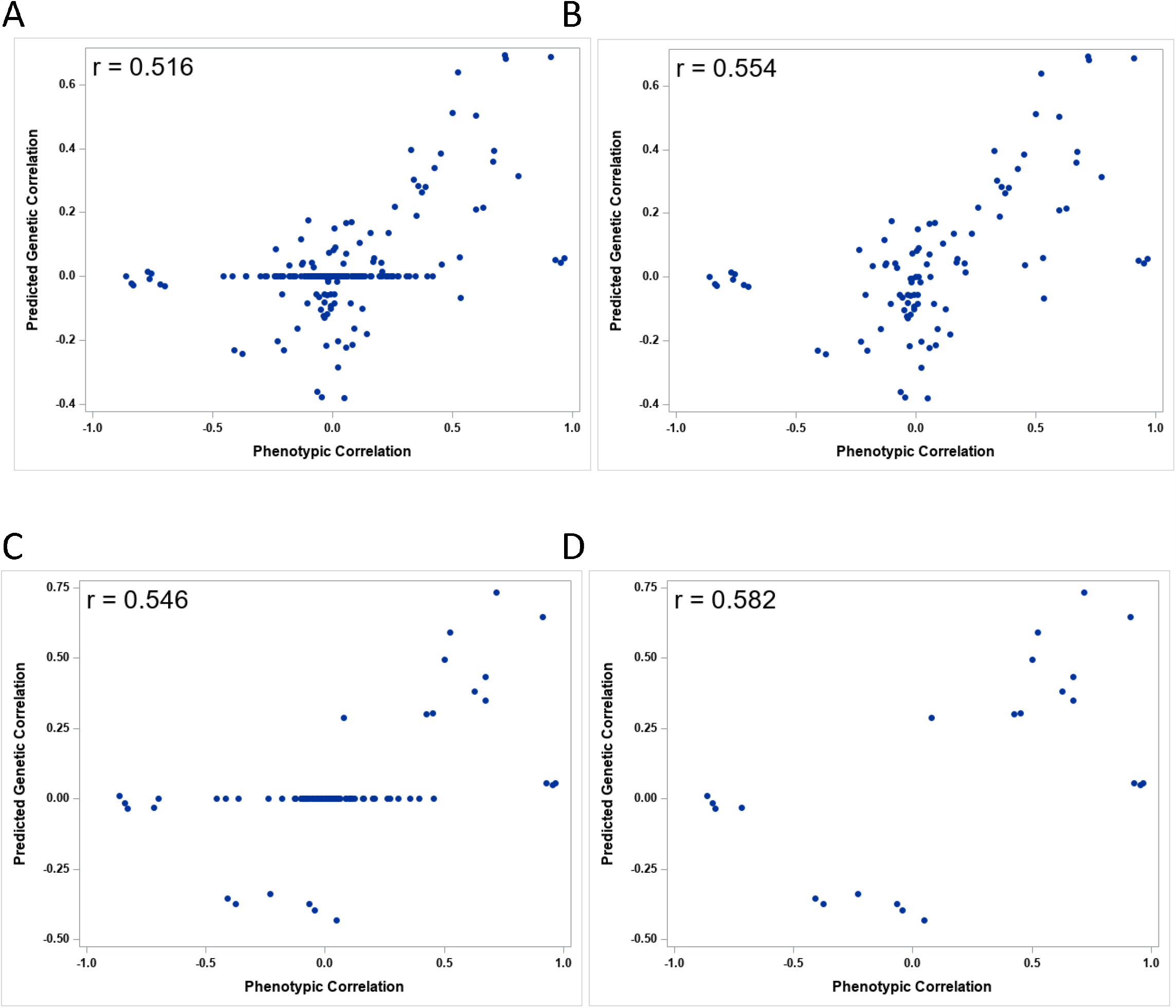
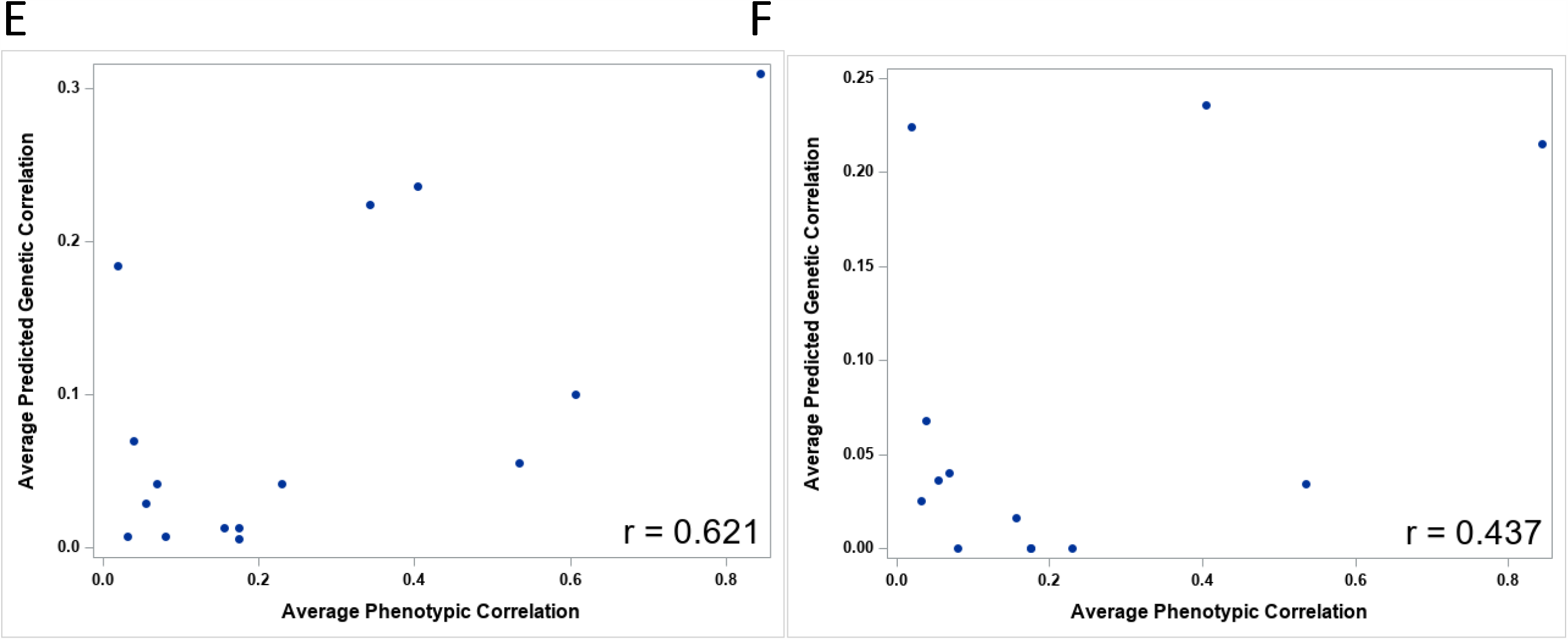
Correlation between predicted genetic and phenotypic correlations. A. – D. Pairwise trait correlations. A and B: all QTLs. C and D: GWS QTLs. A and C: for all trait pairs. B and D: only trait pairs with predicted non-zero genetic correlations. Permutation tests for significance of correlation: A – D. P < 0.001. E. – F. Module average pairwise correlations. E. All QTLs. P = 0.011. F. GWS QTLs. P = 0.066.

Bias, measured as the difference between the predicted genetic correlation and observed phenotypic correlation, increased with the absolute value (magnitude) of phenotypic correlations (Appendix S15), suggesting that for pairs of traits with large phenotypic correlations unidentified QTLs may contribute to those correlations. However, bias was uncorrelated with either average total percent of variance explained or total relative homozygous effect (Appendix S15, Appendix S16), which are indicators of the completeness of QTL identification.

### QTL overlap in other selfing-syndrome species

Our results suggest that trait change within modules may have been correlated in the evolution of the selfing syndrome of *I. lacunosa*, but that between-module correlation likely did not play a major role in this trait divergence. To determine the extent to which this was the case in other species exhibiting the selfing syndrome, we applied our measure of QTL overlap to published QTL studies of the selfing syndrome. For this analysis, we first grouped our traits into modules based on anatomy: floral morphology, inflorescence, phenology, pollen, and nectar (Table 1). Although the genetic architecture of pollination syndromes likely varies between systems (Smith, 2015), anatomical groupings offer the advantage that they can be applied broadly across systems even when other approaches to grouping traits are not available.

Generally, patterns exhibited by the anatomical modules in our study are similar to those for the cluster modules, which supports the usefulness of anatomical groupings when other approaches are unavailable. Average trait correlations are substantially higher within modules than between modules (Appendix S17). QTL overlap was substantially and significantly higher within modules than between modules for both GWS and all QTLs (Appendix S18). The randomization test indicated that there is significantly more QTL overlap within modules than expected by random placement of QTLs (Appendix S19), although this test could not be performed for some modules because there were QTLs for only one trait. The correlation across module pairs between average phenotypic correlation and average QTL overlap was moderate (0.506) and significant for all QTLs, but weak (0.207) and non-significant for GWS QTLs (Appendix S20).

For comparison, we identified nine published QTL mapping studies of the selfing syndrome. They spanned a broad range of angiosperm model systems: three studies in *Mimulus* (Lin and Ritland, 1997; Fishman et al., 2002, 2014), two from *Solanum* (Bernacchi and Tanksley, 1997; Georgiady et al., 2002), three in *Capsella* (Sicard et al., 2011; Slotte et al., 2012; Woźniak et al., 2020), and one from *Leptosiphon* (Goodwillie et al., 2006). We grouped traits from the previous publications into these same anatomical modules. Three studies identified QTLs only for floral morphological traits, while six also included some combination of inflorescence traits, reproductive traits, phenology traits, or vegetative traits (Table 4, Appendix S21). We found similar degrees of overlap in the same and related species even though the studies include spanned a wide range of time and methods.

**Table 4.**
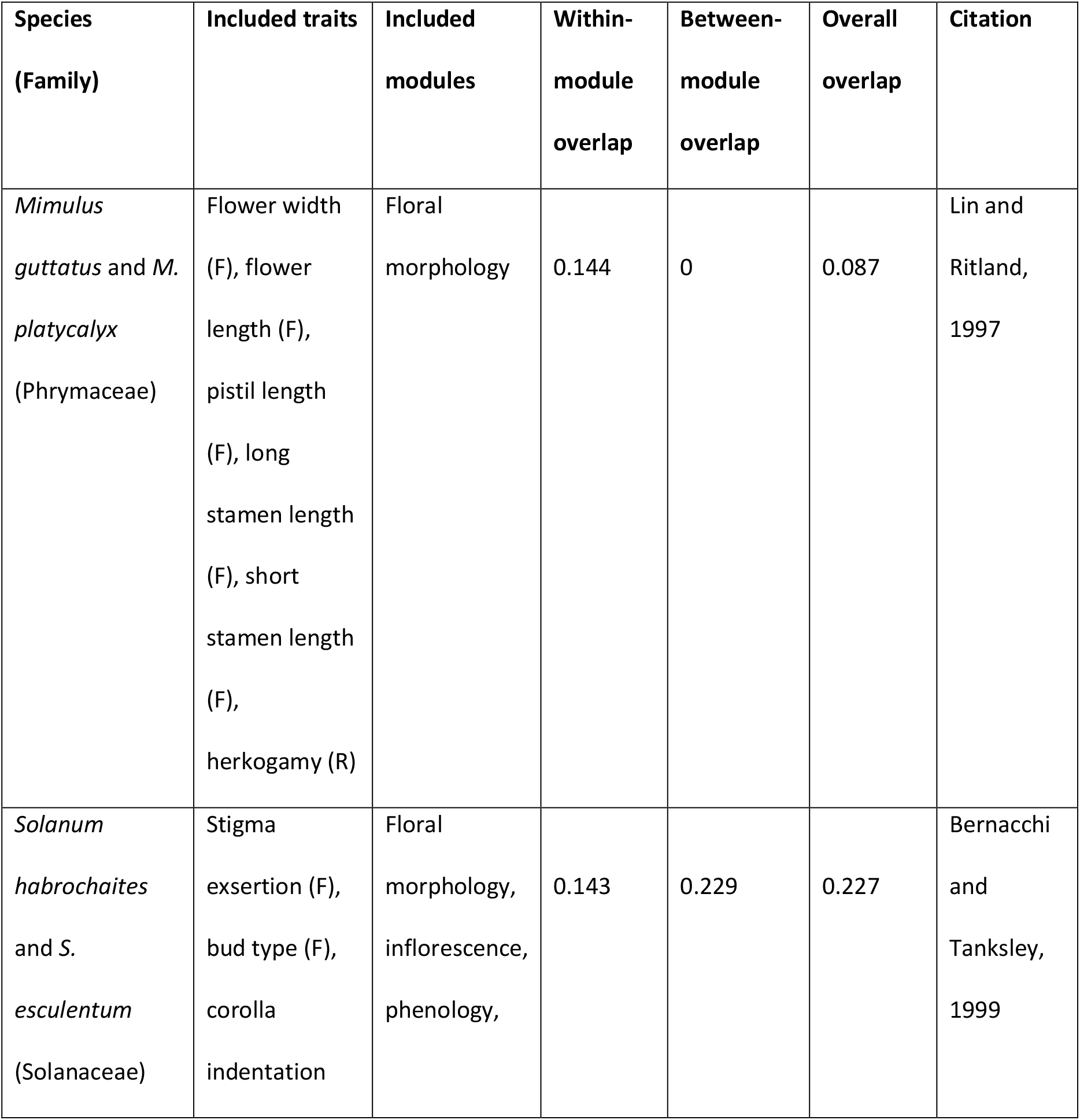
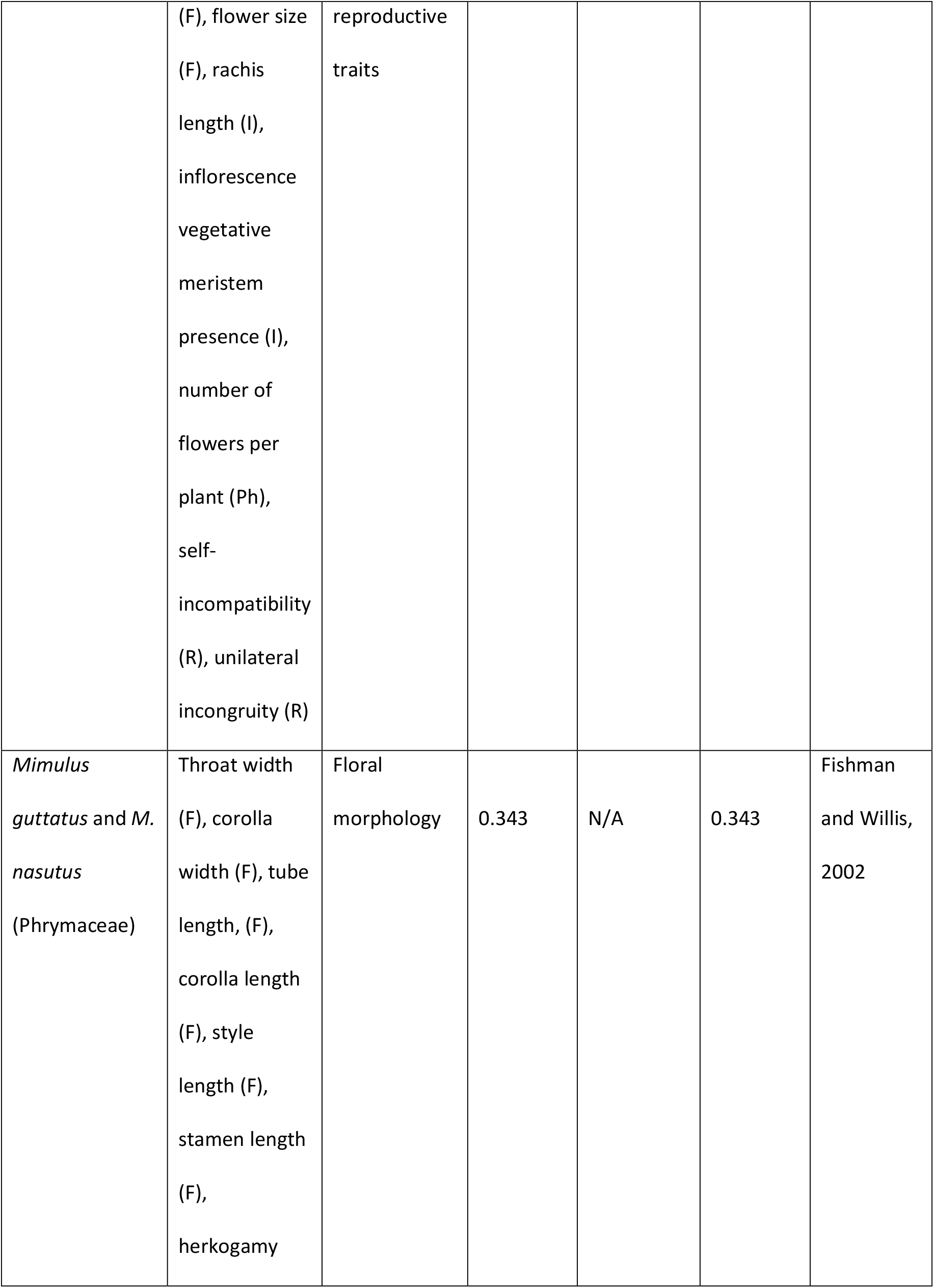
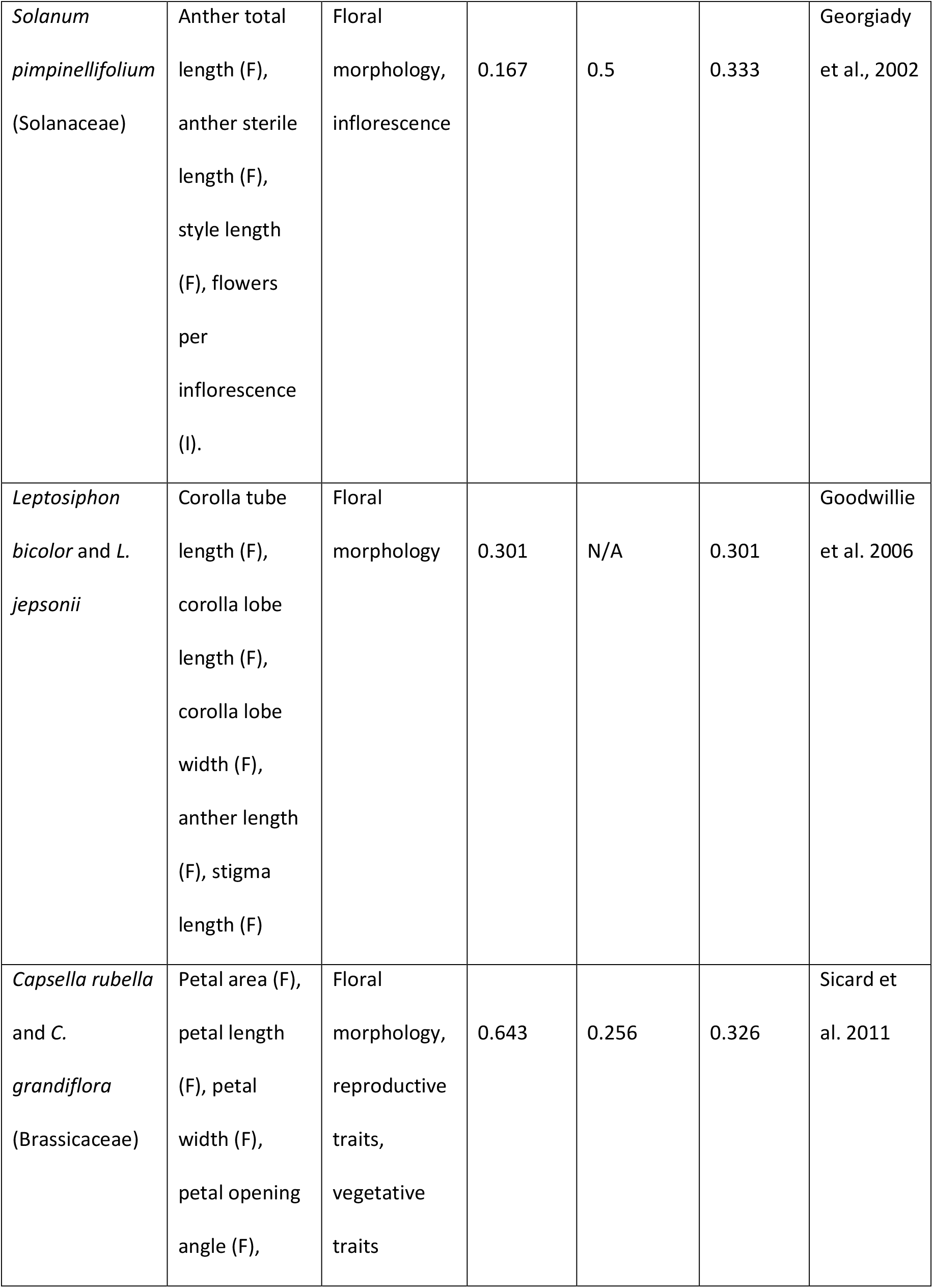
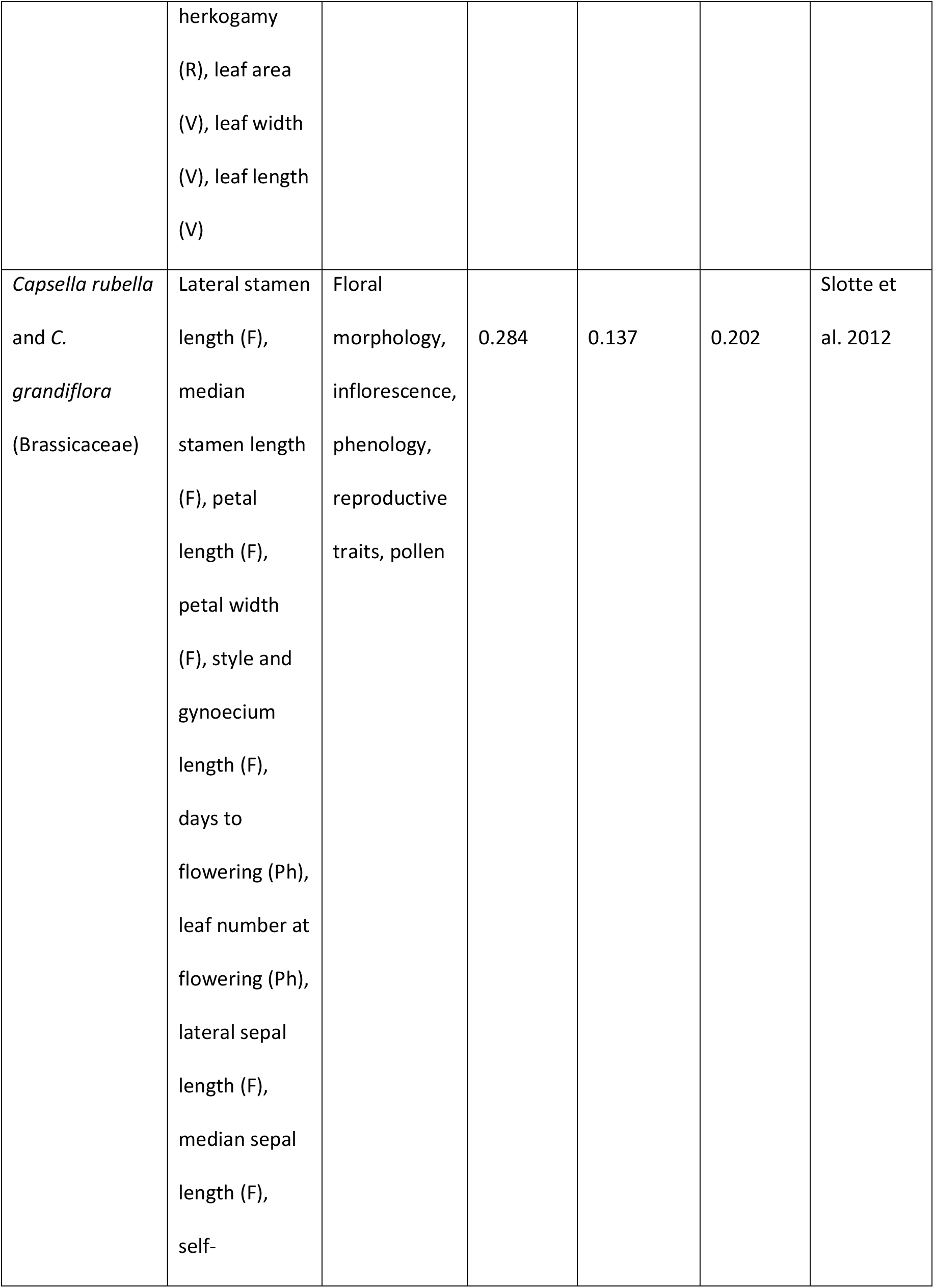
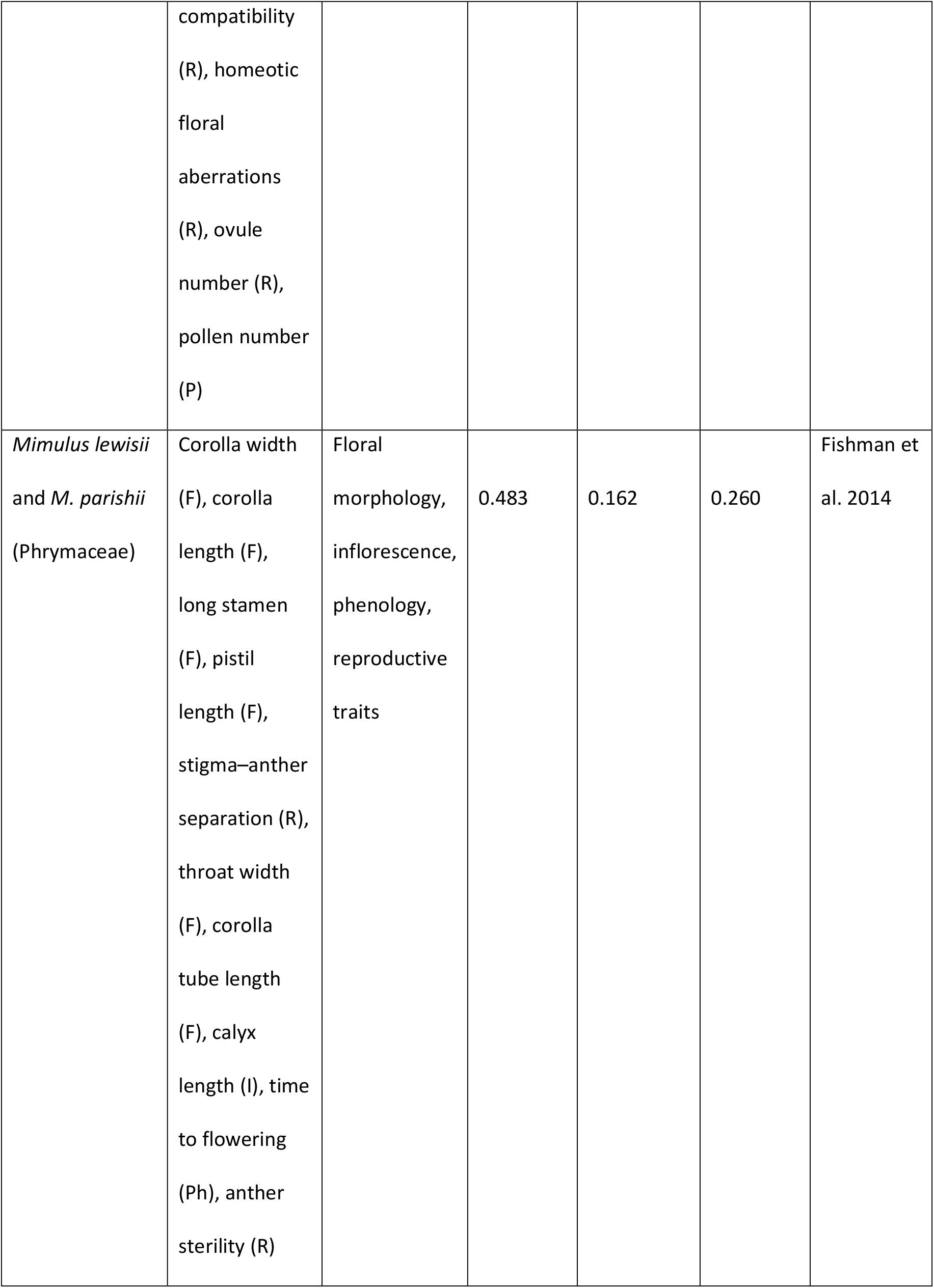
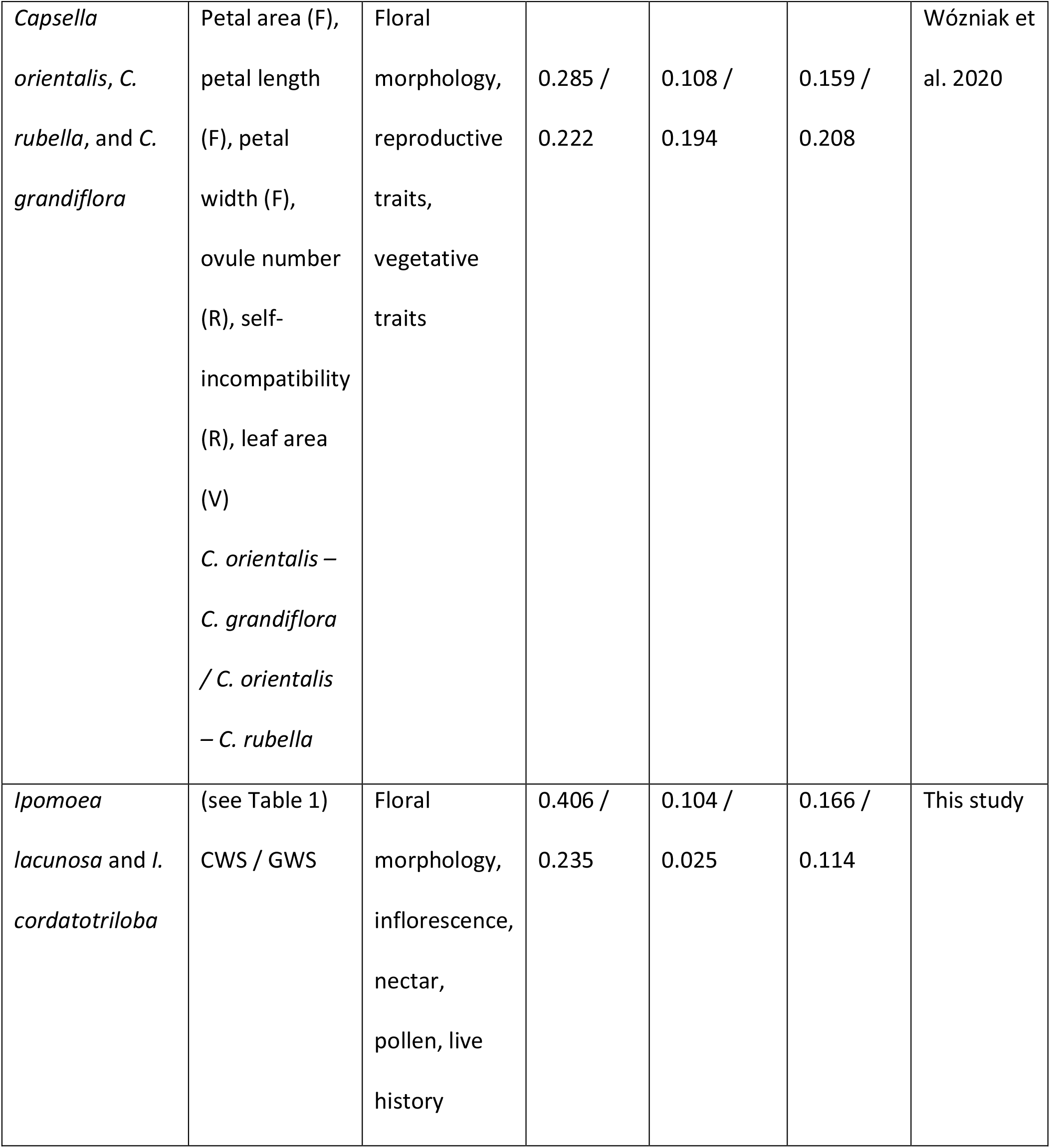
Studies included in our quantitative review of QTL overlap, using anatomical modules. Modules abbreviations are F (floral morphology), C (color traits), I (inflorescence traits), Ph (phenology traits including growth rate and flowering time), N (nectar), Po (pollen), R (reproductive traits such as compatibility), and V (vegetative traits distinct from timing, such as leaf size).

Between-module overlap was lower (mean 0.180) than within-module overlap (mean 0.295) for all systems where it could be calculated except *Solanum*, in which both studies found higher between-than within-module overlap (between: 0.229, 0.5; within: 0.143, 0.167; Bernacchi and Tanksley 1997; Georgiady et al. 2002). However, the level of both within- and between-module overlap varied widely from very high overlap in one *Capsella* study (within: 0.643, between: 0.256; (Sicard et al., 2011)) to fairly low overlap in *Solanum* (within: 0.143-0.167; between: 0-0.5; (Bernacchi and Tanksley, 1997; Georgiady et al., 2002). This is consistent with highly variable importance of genetic correlations in the evolution of different species’ selfing syndromes.

The pattern within floral morphology also reflects this diversity. Floral morphology was examined in the greatest number of studies, and both numbers of QTLs and degrees of overlap varied considerably across systems. *Mimulus* exhibits variable overlap among many widely scattered QTLs (0.144, 0.343, 0.366; (Lin and Ritland, 1997; Fishman, Kelly and Willis, 2002; Fishman *et al*., 2014). In *Solanum*, the degree of overlap and number of regions in play are lower (0.234, 0.167; (Bernacchi and Tanksley, 1997; Georgiady et al., 2002). In *Leptosiphon*, overlap is quite high (0.301; (Goodwillie et al., 2006), and studies in *Capsella* find high concentration of floral morphological traits into few regions (0.286, 277, 0.369, 0.367; (Sicard et al., 2011; Slotte et al., 2012; Woźniak et al., 2020). Therefore the pattern we observed in *Ipomoea lacunosa* floral traits, of numerous loci of small effect with high overlap, is common but by no means universal, and the floral reductions of the selfing syndrome can evolve from a variety of underlying genetic architectures.

Finally, we identified considerable variation in which pairs of modules showed the strongest overlap. In *Capsella*, vegetative traits frequently showed high overlap with floral morphological and reproductive traits (Sicard et al., 2011; Woźniak et al., 2020), which appears to be unusual in flowering plants (Ashman and Majetic, 2006; Feng et al., 2019). In *Mimulus* and *Solanum*, floral morphological traits and inflorescence traits overlapped more (Georgiady et al., 2002; Fishman et al., 2014). In *Ipomoea*, the strongest overlaps were between nectar volume and floral morphology; although this has not been examined in other selfing-syndrome systems, overlapping QTLs have been observed for floral morphology and nectar traits in *Penstemon, Petunia*, and *Mimulus* (reviewed in (Smith, 2015)). Overall, *Ipomoea* fell on the low end of QTL overlap both within and between modules. This study is the first to quantify the diversity of genetic architectures underlying convergently evolved selfing syndromes, and indicates a plethora of possible routes to the same pollination syndrome.

## DISCUSSION

### Architecture of evolutionary divergence

Although the selfing syndrome has evolved repeatedly in angiosperms, the role of correlated evolution, the relative contributions of selection and drift, and what agents of selection act all remain unclear. *Ipomoea lacunosa* is an effective system to explore these problems because selection is implicated in the divergence of at least a subset of its selfing-syndrome traits (Duncan and Rausher, 2013a, 2020; Rifkin, Liao, et al., 2019). *Ipomoea lacunosa* and its mixed-mating sister species *I. cordatotriloba* hybridize in nature, with substantial asymmetric introgression from *I. lacunosa* into *I. cordatotriloba* (Rifkin, Castillo, et al., 2019; average proportion admixture from I. lacunosa > 0.5). Nevertheless, *I. cordatotriloba* sympatric with *I. lacunosa* retain considerable, but not complete, genetic and phenotypic divergence in selfing-syndrome traits, which suggests ongoing divergent selection on these traits. Consequently, the genetic architecture of divergence described here does not reflect the full range of divergence between allopatric populations of the two species. Nevertheless, our results provide insight into the degree to which different components of the selfing syndrome have evolved independently.

If the selfing syndrome of *I. lacunosa* results largely from correlated evolution, we predict high QTL overlap and strongly genetically linked traits. If, on the other hand, selfing-syndrome traits evolved through independent selection on independent traits, we predict low overlap. In our mapping population, both F2 correlation and QTL overlap patterns revealed distinct modules that largely overlap anatomical categories. Traits within modules are moderately to highly correlated and exhibit moderate to high QTL overlap, and within-module trait correlations are overwhelmingly positive. These patterns suggest correlated divergence of traits within modules. By contrast, between-module trait correlations are generally low, as is QTL overlap between traits from different modules. Together, these findings indicate that modules were able to diverge independently, possibly at different times and due to different selective pressures.

A consistent aspect of the selfing syndrome across many species is a reduction in multiple floral characters, including size, pollen production, and nectar production (Sicard and Lenhard, 2011). This can be explained either through a general pattern of high genetic correlation among floral traits, or by independent selection on independent traits. For example, the evolution of reduced floral traits could result from selection to shorten development time, leading to a reduced number of cell divisions (Krizek and Anderson, 2013) that affects all floral traits similarly. In this situation, selection acting directly on only a subset of floral traits could cause a correlated reduction of all floral traits. In *I. lacunosa*, we identified at least two genetically distinct floral modules: one consisting of size traits and probably nectar production, and one consisting of pollen traits. Previous experiments have demonstrated that natural selection contributed to floral size divergence (Rifkin, Liao, et al., 2019), so reductions in floral dimensions likely evolved in a correlated fashion, but the genetic independence of floral dimensions and pollen characters indicates that reduction in pollen production was not a correlated response to selection for floral size reduction. Whether reduced pollen production was caused by directional selection or by reduced purifying selection coupled with genetic drift is unclear; our previous study failed to detect any evidence for selection having favored reduce pollen production in *I. lacunosa*, but evidence from *Arabidopsis thaliana*, another selfing species, implicates selection in pollen reduction (Rifkin, Liao, et al., 2019; Tsuchimatsu et al., 2020).

Although a correlated response to selection can explain the evolution of the floral traits in module 1, the reduction in style length may be an exception. Of the 10 QTL we identified for this trait, six were contra-directional (Appendix S13). However, because only one of these overlapped any other floral QTLs, it seems unlikely that *I. lacunosa* alleles at these loci were fixed because they had advantageous pleiotropic effects on other floral characters. Instead, it seems possible that genetic drift played a substantial role in style reduction despite any correlated response to selection on other floral traits. An alternative explanation that we cannot rule out, however, is that the contra-directional QTL alleles fixed in *I. lacunosa* represent compensatory corrections for correlated responses dragging style length beyond its optimum.

Between-module correlations were generally weaker than within-module correlations, but may also play a role in divergence. For example, cyme length is also diverged between species, but a role for selection has not been identified (Rifkin, Liao, et al., 2019). Although in a distinct module, cyme length is somewhat correlated with floral morphology. Therefore, indirect selection through shared loci that affect floral traits may have caused cyme lengths to diverge in the absence of selection.

In addition to reductions in floral traits, highly selfing species often exhibit changes in vegetative traits, including early growth rate, perhaps because highly selfing species are often found in marginal habitats where rapid growth is at a premium (Snell and Aarssen, 2005). We have previously shown that changes in these types of traits have occurred in *I. lacunosa* (Rifkin, Liao, et al., 2019). Here we have demonstrated that early growth traits are independent from floral traits and constitute two genetically distinct evolutionary modules: one reflecting the rate of internode elongation, the other reflecting general plant size, as reflected in total plant height and number of leaves at day 21, as well as the timing of leaf production. If selection was responsible for these changes, it acted independently on at least two aspects of early growth.

The overall picture that emerges from our study is that the *I. lacunosa* selfing syndrome has evolved piecemeal through changes in at least five distinct groups of traits. Within these modules, correlated responses to selection probably account for much of the change, although in some instances (e.g. evolution of shorter styles) drift may have also played a role. Our interpretation of correlated selection assumes that QTL overlap is indicative of pleiotropy of a single underlying causal variant. Instead, the overlap may represent chance proximity of two different causal variants that were fixed sequentially. Assuming that mutations in many genes scattered throughout the genome could affect a particular trait, it seems unlikely that different variants affecting two traits would by chance be in physical proximity, and likely that average overlap between QTLs affecting those two traits would be low. Our simulations of random placement of QTLs bear this out for within-module trait pairs, and show that the actual overlap is substantially higher than expected by chance placement. These results, as well as evidence of within-module pleiotropy in other systems (Smith, 2015), lend support to the hypothesis that much of the observed QTL overlap is due to pleiotropy.

The low degree of between-module correlation also allows inferences about which possible selective pressures may have acted. Different selective explanations for the selfing syndrome depend on different degrees of trait correlation (Sicard and Lenhard, 2011). Two proposed explanations depend on high correlation: that selection to self-pollinate efficiently could lead to general reductions in floral traits, or that selfing plants colonizing marginal environments experience selective pressure to mature and reproduce faster in these marginal environments, leading to smaller flowers. Other possible explanations impose no such requirement: the selfing syndrome may result from reallocation of resources from pollinator attraction and pollen export to other functions that increase fitness, or selfing plants may reduce floral traits to avoid florivory by insects. The high degree of independence we found among *I. lacunosa*’s selfing-syndrome traits is more consistent with these explanations, which allow traits to evolve independently rather than depending on correlation. Indeed, *I. cordatotriloba* experiences considerable damage from florivorous insects (Sowell and Wolfe, 2010).

### Consistency of approaches

Evolutionary analyses of correlated suites of characters typically rely on inferences based on genetic correlation matrices (e.g. (Delph et al., 2010)). However, genetic correlations are determined by the extent to which QTLs underlying genetic variation have pleiotropic effects on multiple characters, and QTL studies can also be used to both understand and predict genetic correlations (Kelly, 2009; Saltz et al., 2017), although the consistency of these approaches is seldom evaluated. Our study provides evidence that QTL analyses and estimates of F2 correlations provide consistent pictures of the genetic architecture of evolutionary divergence. In particular, we found that evolutionary modules identified by analysis of F2 trait correlations are consistent with patterns of QTL overlap: there is a high correlation between between- and within-module F2 correlations, and average QTL overlaps between and within modules. Additionally, QTL properties predict genetic correlations that exhibit a moderately high correlation with the F2 correlations. These patterns are detectible even though our QTLs on average explain only about 40 percent of the between-species differences.

One caveat to this conclusion is that we measured F2 phenotypic correlations rather than genetic correlations. However, if environmental correlations are not large or not in a direction opposite to the genetic correlations, phenotypic correlations should reflect the underlying genetic correlations (Falconer and Mackay, 1996). Moreover, empirical analyses generally find concordance between genetic and phenotypic correlations (Cheverud, 1988). We thus believe that the F2 phenotypic correlations we measured likely reflect the underlying genetic correlations.

In our study, we detected 43 QTLs with genome-wide significance for which data was available on the direction of trait differences between species, and an additional 38 with just chromosome-wide significance. Similar behavior between the GWS and CWS QTLs suggests that the CWS QTLs are general rather than artifactual. The correlation between F2 correlations and QTL overlap (Fig. 2) is similar when using all QTL rather than just GWS QTL. If the CWS QTLs were artifactual, we would expect that an analysis including them would reduce the strength of the correlation. Similarly, the difference in average pairwise QTL overlap between within- and between-module comparisons was similar for analyses using all QTL and just GWS QTL. Again, if the CWS QTL were artifactual, we would expect that the difference would be substantially reduced in the former analysis. We also observed that the direction of QTL effects (in same or opposite direction as species difference) was highly biased against contra-directional effects (Table S6), and the proportion of consistent- vs contra-directional effects was not statistically different between GWS and CWS, with both types of QTL exhibiting significant bias toward QTL with consistent-direction effects (P < 0.005 in both cases). Artifactual QTLs should exhibit no bias in the direction of effects. We believe these similarities between results with GWS and CWS QTLs justify treating all QTLs as legitimate.

### Comparative genetic architecture of the selfing syndrome

The selfing syndrome has evolved repeatedly in numerous angiosperm lineages. Our quantitative review places the genetic architecture of the *I. lacunosa* selfing syndrome in a wider context of selfing syndrome evolution. Across the studies we examined, we found QTL overlap to be highly variable, both within and between modules. Floral traits, which have been studied most extensively, show a wide range of variation for within-module overlap. In *Mimulus*, most studies find high QTL overlap within floral traits (Lin and Ritland, 1997; Fishman et al., 2002, 2014), and that floral traits are controlled by pleiotropic QTLs scattered across the genome. Overlap is high, but overall numbers of QTL lower, in *Leptosiphon* and *Capsella* (Goodwillie et al., 2006; Sicard et al., 2011; Slotte et al., 2012; Woźniak et al., 2020), while *Solanum* had low overlap for floral traits and few regions (Bernacchi and Tanksley, 1997; Georgiady et al., 2002). Between-module overlap also varies widely across systems. In general, floral traits are expected to be more interconnected with each other than with vegetative traits (Fenster and Armbruster, 2004; Ashman and Majetic, 2006; Glover et al., 2015; Smith, 2015), and consistent with this, we generally found higher within-than between-module overlaps. However, average between-module overlap varied widely, and indeed, in the genus *Solanum* both flowering time (Bernacchi and Tanksley, 1997) and inflorescence structure (Georgiady et al., 2002) were so strongly correlated with floral traits that between-module correlation was stronger than within-module correlation, and a clustering approach to module identification would likely identify different underlying genetic groupings.

It is unlikely that genome structure alone drives this variation in genetic architecture. The species included in these studies vary in genome size and number of chromosomes, but *Ipomoea* and *Mimulus*, which have very different patterns of QTL overlap, are similar in genome size (*M. guttatus*, 430Mb; *I. lacunosa* and *I. cordatotriloba*, 497Mb and 525Mb respectively) and chromosome number (*M. guttatus*, 2N=28, *I. lacunosa* and *I. cordatotriloba*, 2N=30) (Duncan and Rausher, 2013b; Institute, 2017). The variation we noted in genetic architecture is also not explained by the number or type of traits included: studies in the same systems are broadly similar across time, and there was extensive variation both in studies that included floral and phenology traits and in studies that only included floral traits. This variation in genetic architecture suggests that selective causes for the selfing syndrome may vary across systems: in *Ipomoea* we can reject explanations for selfing-syndrome evolution that depend heavily on genetic correlation, but they likely occur in other systems.

### Conclusions and future directions

The research we describe here identifies two areas of high priority for future research in determining how and why the selfing syndrome generally evolves.

First, much remains unknown about the relative importance of the possible selective causes in selfing syndrome evolution across angiosperms. It is possible to make specific evolutionary predictions based on these selective explanations, but they must be tested with empirical data. For example, nectar volume is unlikely to affect the efficiency of selfing, but may influence insect predators. Similarly, the selfing syndrome can only evolve in response to selection for faster global development or as a byproduct of selection for more efficient selfing in species with strong correlations between traits. We predict that species with greater independence are more likely to evolve the selfing syndrome through resource allocation or evading florivory. Field tests of selection in species with known genetic architecture and natural history could evaluate these predictions.

The variation in genetic architecture also invites questions about how genetic architecture itself evolves (Hansen, 2013). We generally found consistent genetic architecture within the same systems, but one study has identified decreased floral trait integration in selfing species compared to outcrossing relatives, so this pattern may be weaker in selfing species than in other pollination syndrome shifts (Anderson and Busch, 2006). Further understanding the relationship between genetic architecture and the transition to selfing is another priority moving forward in efforts to understand the relationship between selection, drift, and genetic architecture in the evolution of the selfing syndrome.

## Supporting information

Rifkin_Appendix_S9_Cor_plot_scatterplots_and_distributions

Rifkin_Appendix_S10_cluster_diagrams

Rifkin_Appendix_S11_PVE_and_RHE

Rifkin_Appendix_S12_Average_module_PVE_and_RHE

Rifkin_Appendix_S13_QTL_directions

Rifkin_Appendix_S14_predicted_correlation_correlations

Rifkin_Appendix_S15_bias_vs_PVE_RHE

Rifkin_Appendix_S16_bias_vs_PVE_RHE_GWS_only

Rifkin_Appendix_S17_anatomical_module_correlations

Rifkin_Appendix_S18_QTL_overlap_anatomical_modules

Rifkin_Appendix_S19_QTL_localization

Rifkin_Appendix_S20_predicted_correlation_correlations_anatomical

Rifkin_Appendix_S21_Quant_review_supp_table

Rifkin_Appendix_S1_Measurement_methods

Rifkin_Appendix_S2_ddRADseq_library_construction_and_sequencing

Rifkin_Appendix_S3_Linkage map construction

Rifkin_Appendix_S4_QTL_mapping

Rifkin_Appendix_S5_QTL_positions

Rifkin_Appendix_S6_Genome_annotation

Rifkin_Appendix_S7_Predicting_Genetic_Correlations_from_QTL_properties

Rifkin_Appendix_S8_Trait_correlations

## ACKNOWLEDGMENTS

We thank Tanya Duncan for support in identifying populations and developing the mapping population, and Shu-Mei Chang and Stephen DiMaria for assistance with pollen measurements. Funding was provided by NSF grant DEB 1542387 to MDR, by NSF DDIG grant DEB 1501954 to JLR.

## AUTHOR CONTRIBUTIONS

JLR and MDR conceived the design, analyzed the data, and wrote the manuscript. JLR generated, grew, and phenotyped the mapping population, constructed the genetic libraries, and performed the QTL mapping and quantitative review. MDR performed the cluster analyses, comparison of predicted genetic correlations with QTL overlap, and analysis of QTL clustering. GC performed the genome annotation and calculated gene densities.

## DATA AVAILABILITY STATEMENT

Raw sequencing reads from the F2 mapping population will be available on the SRA (PRJNA691909, embargoed until publication). Scripts are available on Github (https://github.com/joannarifkin/Ipomoea_QTL).

## WORKS CITED

Anderson, I., and J. Busch. 2006. Pollinator-mediated selection weakens floral integration in self- compatible taxa of Leavenworthia (Brassicaceae). American Journal of Botany 93: 860–867.

Armbruster, W. S., C. Pélabon, G. H. Bolstad, and T. F. Hansen. 2014. Integrated phenotypes: Understanding trait covariation in plants and animals. Philosophical Transactions of the Royal Society B: Biological Sciences 369.

Ashman, T.-L., and C. J. Majetic. 2006. Genetic constraints on floral evolution: a review and evaluation of patterns. Heredity 96: 343–352.

Barrett, S. C. 2002. The evolution of plant sexual diversity. Nature Reviews Genetics 3: 274–84.

Bernacchi, D., and S. Tanksley. 1997. An interspecific backcross of Lycopersicon esculentum x L. hirsutum: linkage analysis and a QTL study of sexual compatibility factors and floral traits. Genetics 14850: 861–877.

Brandon, R. N. 1999. The units of selection revisited: the modules of selection. Biology and Philosophy 14: 167–180.

Broman, K. W., D. M. Gatti, P. Simecek, N. A. Furlotte, P. Prins, S. Sen, B. S. Yandell, and G. A. Churchill. 2019. R/qtl2: Software for mapping quantitative trait loci with high-dimensional data and multiparent populations. Genetics 211: 495–502.

Cheverud, J. M. 1988. A comparison of genetic and phenotypic correlations. Evolution 42: 958–968.

Delph, L. F., A. M. Arntz, C. Scotti-Saintagne, and I. Scotti. 2010. The genomic architecture of sexual dimorphism in the dioecious plant Silene latifolia. Evolution 64: 2873–2886.

Duncan, T. M., and M. D. Rausher. 2013a. Evolution of the selfing syndrome in Ipomoea. Frontiers in Plant Science 4: 1–8.

Duncan, T. M., and M. D. Rausher. 2013b. Morphological and genetic differentiation and reproductive isolation among closely related taxa in the Ipomoea series Batatas. American Journal of Botany 100: 2383–2193.

Duncan, T. M., and M. D. Rausher. 2020. Selection favors loss of floral pigmentation in a highly selfing morning glory. PLoS ONE 15: 1–18.

Falconer, D. S., and T. F. C. Mackay. 1996. Introduction to quantitative genetics. Longman Group, Essex.

Feng, C., C. Feng, L. Yang, M. Kang, and M. D. Rausher. 2019. Genetic architecture of quantitative flower and leaf traits in a pair of sympatric sister species of Primulina. Heredity 122: 864–876.

Fenster, C., and W. Armbruster. 2004. Pollination syndromes and floral specialization. Annual Review of Ecology, Evolution, and Systematics 35: 375–403.

Fishman, L., P. M. Beardsley, A. Stathos, C. F. Williams, and J. P. Hill. 2014. The genetic architecture of traits associated with the evolution of self-pollination in Mimulus. The New Phytologist 205: 907–917.

Fishman, L., A. J. Kelly, and J. H. Willis. 2002. Minor quantitative trait loci underlie floral traits associated with mating system divergence in Mimulus. Evolution 56: 2138–55.

Gardner, K. M., and R. G. Latta. 2007. Shared quantitative trait loci underlying the genetic correlation between continuous traits. Molecular Ecology 16: 4195–4209.

Georgiady, M., R. Whitkus, and E. Lord. 2002. Genetic analysis of traits distinguishing outcrossing and self-pollinating forms of currant tomato, Lycopersicon pimpinellifolium (Jusl.) Mill. Genetics 344: 333–344.

Glover, B. J., C. A. Airoldi, S. F. Brockington, M. Fernández-Mazuecos, C. Martínez-Pérez, G. Mellers, E. Moyroud, and L. Taylor. 2015. How have advances in comparative floral development influenced our understanding of floral evolution? International Journal of Plant Sciences 176: 307–323.

Goodwillie, C., C. Ritland, and K. Ritland. 2006. The genetic basis of floral traits associated with mating system evolution in Leptosiphon (Polemoniaceae): an analysis of quantitative trait loci. Evolution 60: 491–504.

Hansen, T. E. 2013. The evolution of genetic architecture. Annual Review of Ecology, Evolution, and Sytematics 37: 123–157.

Harrell Jr, F. E., with contributions from Charles Dupont, and many others. 2016. Hmisc: Harrell Miscellaneous.

Institute, D. J. G. 2017. Mimulus Genome Project. Institute, DoE Joint Genome. Website http://genome.jgi.doe.gov/mimulus/mimulus.home.html.

Kelly, J. K. 2009. Connecting QTLs to the G-matrix of evolutionary quantitative genetics. Evolution 63: 813–825.

Kostyun, J. L., M. J. S. S. Gibson, C. M. King, and L. C. Moyle. 2019. A simple genetic architecture and low constraint allows rapid floral evolution in a diverse and recently radiating plant genus. New Phytologist 1: nph.15844.

Krizek, B. A., and J. T. Anderson. 2013. Control of flower size. Journal of Experimental Botany 64: 1427–1437.

Lande, R., and S. J. Arnold. 1983. The measurement of selection on correlated characters. Evolution 37: 1210–1226.

Lendvai, G., and D. A. Levin. 2003. Rapid response to artificial selection on flower size in Phlox. Heredity 90: 336–342.

Lin, J., and K. Ritland. 1997. Quantitative trait loci differentiating the outbreeding Mimulus guttatus from the inbreeding M. platycalyx. Genetics 146: 1115–1121.

Nakajima, G. 1963. Karyotype of genus Ipomoea. Cytologia 28: 351–359.

Ornduff, R. 1969. Reproductive biology in relation to systematics. Taxon 18: 121–133.

Peterson, B. K., J. N. Weber, E. H. Kay, H. S. Fisher, and H. E. Hoekstra. 2012. Double digest RADseq: an inexpensive method for de novo SNP discovery and genotyping in model and non-model species. PloS one 7: e37135.

Rastas, P. 2020. Lep-Anchor: automated construction of linkage map anchored haploid genomes. Bioinformatics (Oxford, England) 36: 2359–2364.

Rastas, P., L. Paulin, I. Hanski, R. Lehtonen, P. Auvinen, and M. Brudno. 2013. Lep-MAP: Fast and accurate linkage map construction for large SNP datasets. Bioinformatics 29: 3128–3134.

Rifkin, J. L., A. S. Castillo, I. T. Liao, and M. D. Rausher. 2019. Gene flow, divergent selection and resistance to introgression in two species of morning glories (Ipomoea). Molecular Ecology 28: 1709–1729.

Rifkin, J. L., I. T. Liao, A. S. Castillo, and M. D. Rausher. 2019. Multiple aspects of the selfing syndrome of the morning glory Ipomoea lacunosa evolved in response to selection: A Qst - Fst comparison. Ecology and Evolution: 7712–7725.

Roff, D. A. 1995. The estimation of genetic correlations from phenotypic correlations: A test of Cheverud’s conjecture. Heredity 74: 481–490.

Salih, H., and D. L. Adelson. 2009. QTL global meta-analysis: Are trait determining genes clustered? BMC Genomics 10: 1–8.

Saltz, J. B., F. C. Hessel, and M. W. Kelly. 2017. Trait Correlations in the Genomics Era. Trends in Ecology and Evolution 32: 279–290.

SAS Instititute. 2013. SAS version 9.4.

Sicard, A., and M. Lenhard. 2011. The selfing syndrome: A model for studying the genetic and evolutionary basis of morphological adaptation in plants. Annals of Botany 107: 1433–43.

Sicard, A., N. Stacey, K. Hermann, J. Dessoly, B. Neuffer, I. Bäurle, and M. Lenhard. 2011. Genetics, evolution, and adaptive significance of the selfing syndrome in the genus Capsella. Plant Cell 23: 3156–3171.

Slotte, T., K. Hazzouri, and D. Stern. 2012. Genetic architecture and adaptive significance of the selfing syndrome in Capsella. Evolution 66: 1360–1374.

Smith, S. D. 2015. Pleiotropy and the evolution of floral integration. New Phytologist 209: 80–85.

Snell, R., and L. W. Aarssen. 2005. Life history traits in selfing versus outcrossing annuals: exploring the ‘time-limitation’ hypothesis for the fitness benefit of self-pollination. BMC Ecology 5: 2.

Sowell, D. D. R., and L. L. M. Wolfe. 2010. Pattern and consequences of floral herbivory in four sympatric Ipomoea species. The American Midland Naturalist 163: 173–185.

Stern, D. L. 2013. The genetic causes of convergent evolution. Nature Reviews Genetics 14: 751–64.

Tsuchimatsu, T., H. Kakui, M. Yamazaki, C. Marona, H. Tsutsui, A. Hedhly, D. Meng, et al. 2020. Adaptive reduction of male gamete number in the selfing plant Arabidopsis thaliana. Nature Communications 11: 2885.

USDA, and NRCS. 2017. The PLANTS Database. National Plant Data Team, Greensboro NC USA.

Wagner, G. P., and L. Altenberg. 1996. Complex adaptations and the evolution of evolvability. Evolution 50: 967–976.

Waitt, D. E., and D. A. Levin. 1998. Genetic and phenotypic correlations in plants: A botanical test of Cheverud’s conjecture. Heredity 80: 310–319.

Wessinger, C. A., and L. C. Hileman. 2016. Accessibility, constraint, and repetition in adaptive floral evolution. Developmental biology: 1–9.

Wickham, H., R. François, L. Henry, and K. Müller. 2020. dplyr: A grammar of data manipulation.

Worley, A. C., and S. C. Barrett. 2000. Evolution of floral display in Eichhornia paniculata (Pontederiaceae): direct and correlated responses to selection on flower size and number. Evolution 54: 1533–1545.

Woźniak, N. J., C. Kappel, C. Marona, L. Altschmied, B. Neuffer, and A. Sicard. 2020. A similar genetic architecture underlies the convergent evolution of the selfing syndrome in Capsella. The Plant Cell 32: 935–949.

Woźniak, N. J., and A. Sicard. 2018. Evolvability of flower geometry: Convergence in pollinator-driven morphological evolution of flowers. Seminars in Cell & Developmental Biology 79: 3–15.

