## Supplementary figures and images for "GENETIC ARCHITECTURE OF DIVERGENCE: THE SELFING SYNDROME IN *IPOMOEA LACUNOSA*"

### Rifkin_Appendix_S9_Cor_plot_scatterplots_and_distributions

Correlations between traits

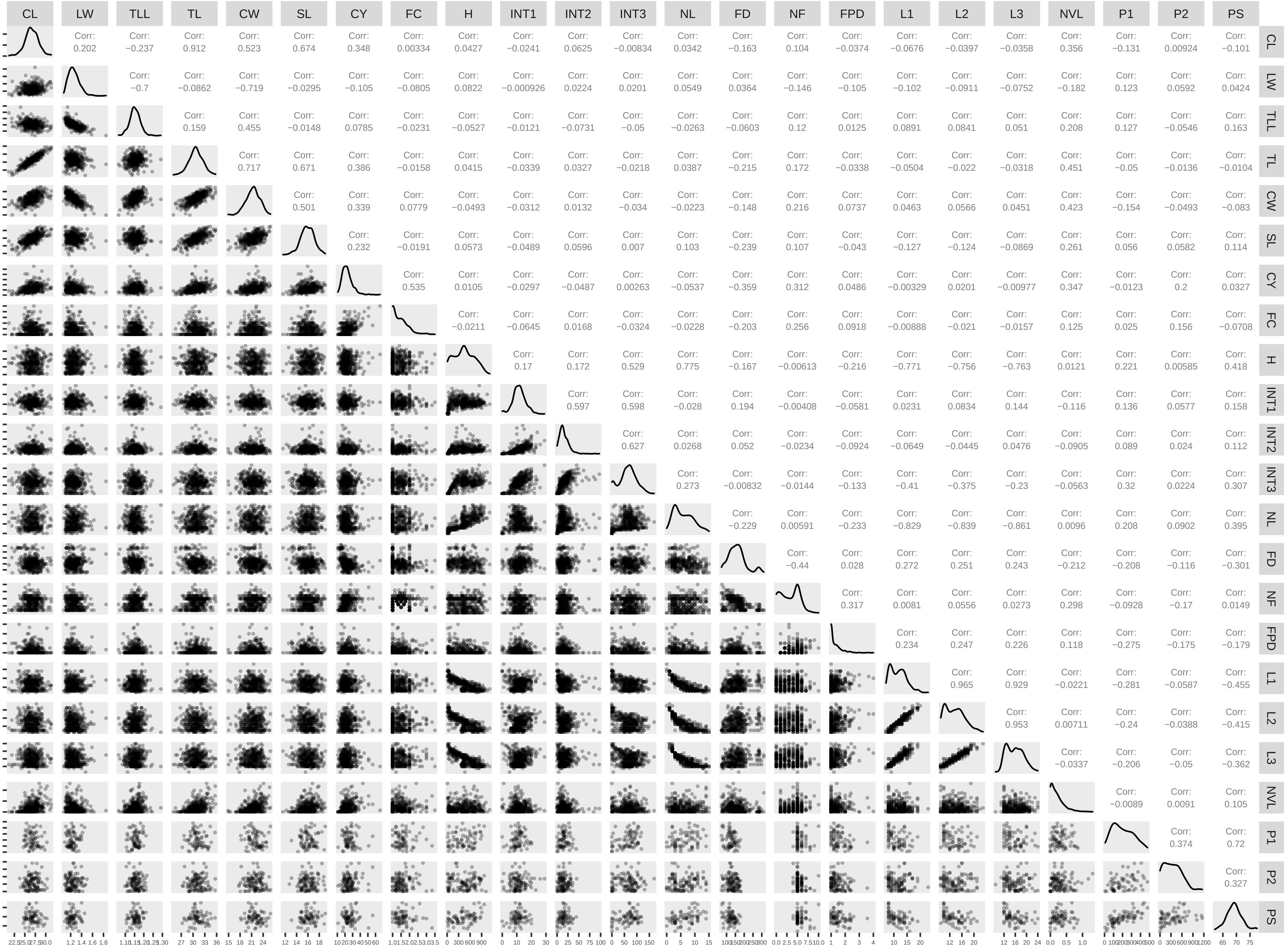

### Rifkin_Appendix_S10_cluster_diagrams

Appendix S9. Cluster diagrams for averaging (A), Ward (B), and McQuitty (C) algorithms.

A

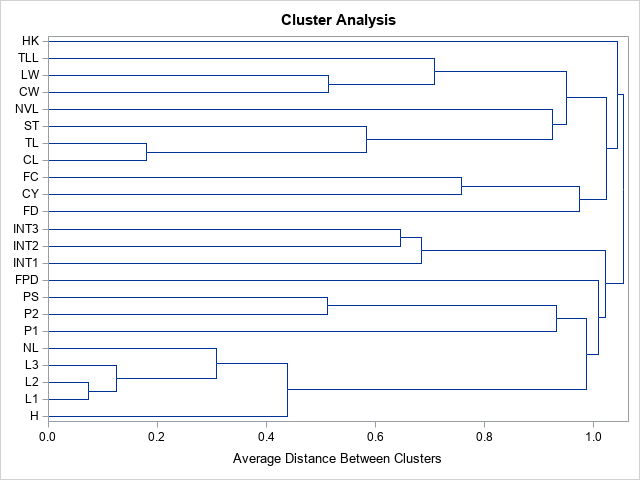

B

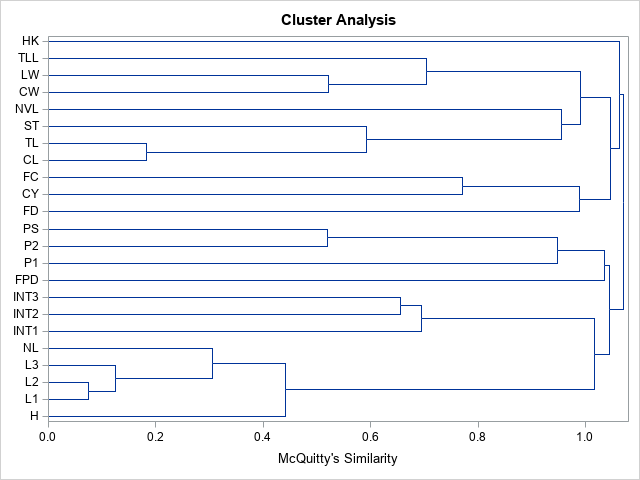

C

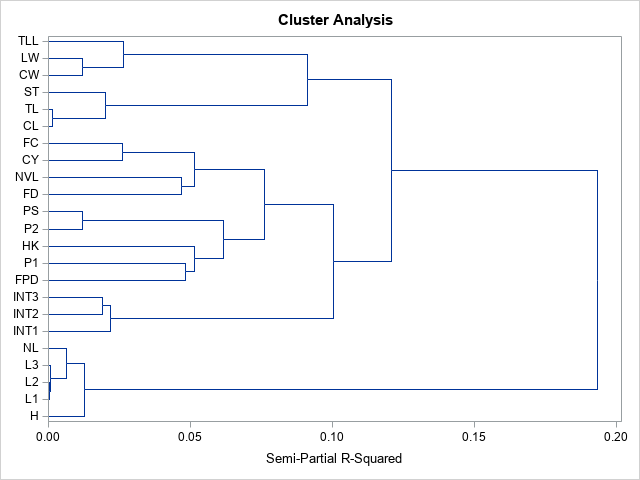
