## Supplementary material for "GENETIC ARCHITECTURE OF DIVERGENCE: THE SELFING SYNDROME IN *IPOMOEA LACUNOSA*": Rifkin_Appendix_S11_PVE_and_RHE

Appendix S10 Total Percent Variance Explained (PVE) and total Relative Homozygous Effect (RHE) for traits.

“ –“ indicates total RHE not calculable because there is no data on parental difference.

| Trait | total PVE | total RHE |
| --- | --- | --- |
| Corolla Length (CL) | 0.540601 | 0.414002 |
| Corolla Tissue Length (TL) | 0.493688 | 0.333482 |
| Corolla Width (CW) | 0.292081 | 0.295691 |
| Corolla Shape 1 (LW) | 0.291875 | 1.172948 |
| Corolla Shape 2 (TLL) | 0.134968 | 0.616183 |
| Style Length (ST) | 0.691264 | 0.646634 |
| Nectar Volume (NVL) | 0.249451 | 0.303531 |
| Flowers or Buds on Cyme (FC) | 0.036516 | -- |
| Cyme Length (CY) | 0.039301 | 0.443856 |
| Flowering Date (FD) | 0.135563 | -- |
| Length of First Internode (INT1) | 0.152847 | 0.449143 |
| Length of Second Internode (INT2) | 0.054232 | 0.453657 |
| Length of Third Internode (INT3) | 0.172119 | 0.695985 |
| Pollen Count 1 (P1) | 0.075849 | 0.241194 |
| Pollen Count 2 (P2) | 0.076201 | 0.437707 |
| Pollen Diameter (PS) | 0.051943 | 0.160742 |
| Flowers Per Day (FPD) | 0.200877 | -- |
| First Leaf Opening (L1) | 0.22649 | -- |
| Second Leaf Opening (L2) | 0.186008 | -- |
| Third Leaf Opening (L3) | 0.210764 | -- |
| Number of Leaves (NL) | 0.2176 | -- |
| Height (H) | 0.034723 | 1.299765 |
| Number of Flowers Open (FPD) | 0.488195 | -- |
