## Supplementary material for "GENETIC ARCHITECTURE OF DIVERGENCE: THE SELFING SYNDROME IN *IPOMOEA LACUNOSA*": Rifkin_Appendix_S12_Average_module_PVE_and_RHE

Appendix S11. Average percent variance explained (PVE) and relative homozygous effect (RHE) and associated standard errors for cluster modules.

| module | Ave PVE | Std Error | Ave RHE | Std Error |
| --- | --- | --- | --- | --- |
| 1 | 0.384847 | 0.068216 | 0.540353 | 0.10991 |
| 2 | 0.07046 | 0.026586 | 0.443856 | NA |
| 3 | 0.126399 | 0.02981 | 0.532928 | 0.066576 |
| 4 | 0.101217 | 0.029186 | 0.279881 | 0.067165 |
| 5 | 0.175117 | 0.031965 | NA | NA |
