## Supplementary material for "GENETIC ARCHITECTURE OF DIVERGENCE: THE SELFING SYNDROME IN *IPOMOEA LACUNOSA*": Rifkin_Appendix_S13_QTL_directions

### Appendix S12 Numbers of QTLs with effects in the direction consistent with direction of trait difference between species (consistent-direction QTLs) and in the opposite direction (contra-directional QTLs) for QTLs of different statistical significance.

Only QTLs for which direction of species difference is known are tabulated. A. Numbers for individual traits. B. Sum over traits. GWS: QTL’s showing genome-wide significance. CWS: QTL’s showing only chromosome-wide significance. Difference in proportions between GWS and CWS: P = 0.120, Fisher exact test. Deviation from expectation of equal numbers of consistent- and contra-directional QTLs: for GWS, χ^2^ = 7.714, df = 1, P = 0.00548; for CWS, χ^2^ = 15.158, df = 1, P = 0.00099.

A.

|  |  |  | GWS |  |  | CWS |  |
| --- | --- | --- | --- | --- | --- | --- | --- |
| Trait |  |  | consistent | contra |  | consistent | contra |
| CL |  |  | 4 | 0 |  | 5 | 0 |
| CW |  |  | 6 | 0 |  | 0 | 0 |
| TL |  |  | 6 | 0 |  | 1 | 0 |
| LW |  |  | 2 | 1 |  | 2 | 1 |
| TLL |  |  | 1 | 0 |  | 1 | 1 |
| SL |  |  | 3 | 3 |  | 1 | 3 |
| CY |  |  | 0 | 0 |  | 1 | 0 |
| FC_binary |  |  | 1 | 0 |  | 1 | 0 |
| FC_continuous | |  | 0 | 0 |  | 0 | 1 |
| H |  |  | 0 | 0 |  | 1 | 0 |
| INT1 |  |  | 0 | 0 |  | 4 | 0 |
| INT2 |  |  | 1 | 0 |  | 0 | 0 |
| INT3 |  |  | 1 | 0 |  | 3 | 0 |
| NL |  |  | 1 | 2 |  | 1 | 0 |
| FD |  |  |  |  |  |  |  |
| NF |  |  |  |  |  |  |  |
| FPD |  |  |  |  |  |  |  |
| EF |  |  | - | - |  | - | - |
| L1 |  |  | 1 | 2 |  | 1 | 0 |
| L2 |  |  | 0 | 2 |  | 1 | 0 |
| L3 |  |  | 1 | 1 |  | 1 | 1 |
| NVL |  |  | 1 | 0 |  | 3 | 1 |
| P1 |  |  | 0 | 0 |  | 2 | 0 |
| P2 |  |  | 0 | 0 |  | 2 | 0 |
| PS |  |  | 1 | 0 |  | 0 | 0 |
| NF |  |  | - | - |  | - | - |

B.

Direction of QTL effect	Proportion

Consistent	Contra	Consistent

GWS	30	11	0.732

CWS	31	8	0.795
