## Supplementary material for "GENETIC ARCHITECTURE OF DIVERGENCE: THE SELFING SYNDROME IN *IPOMOEA LACUNOSA*": Rifkin_Appendix_S14_predicted_correlation_correlations

Appendix S13. The relationship between bias in predicted genetic correlations and observed phenotypic correlations (A-D), between bias and total percent variance explained by QTLs (E-H), and between bias and total relative homozygous effect (I-L).

A. All QTLs, all trait pairs. B. All QTLs, only trait pairs with predicted r ≠ 0. C. GWS QTLs, all trait pairs. D. GWS QTLs, only trait pairs with predicted r ≠ 0. All correlations in A-D significant by permutation test at P < 0.001.

A	B

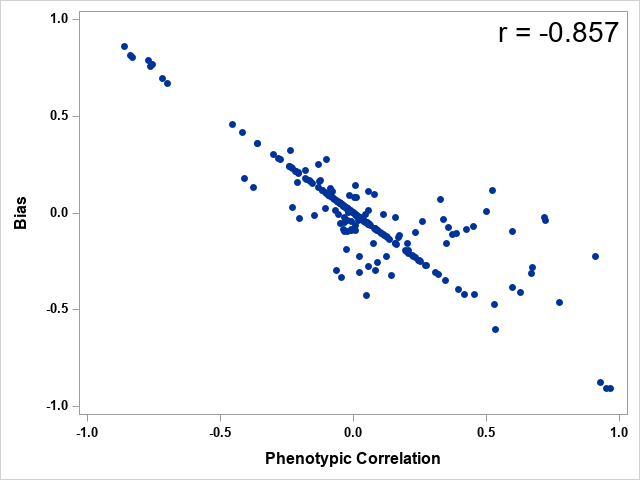

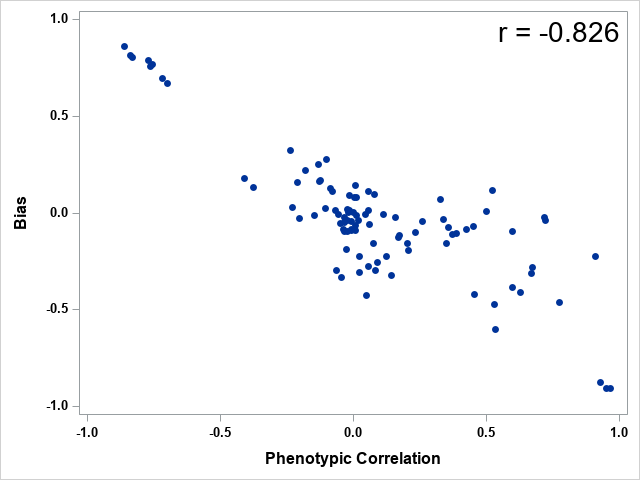

C	D

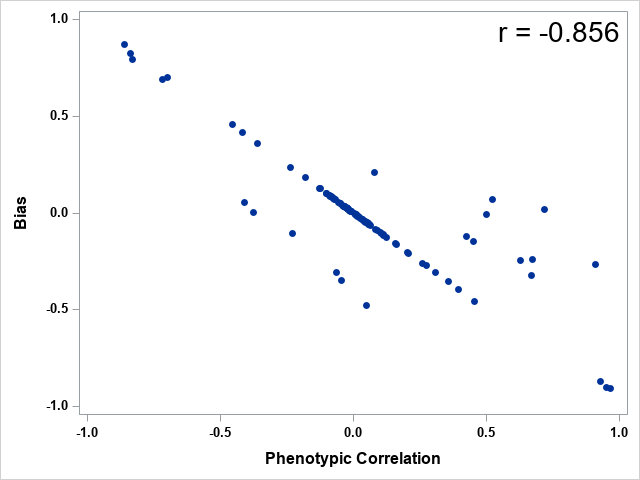

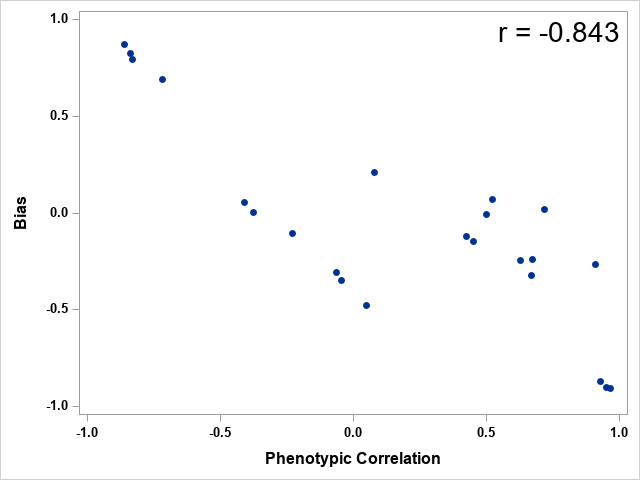
