## Supplementary material for "GENETIC ARCHITECTURE OF DIVERGENCE: THE SELFING SYNDROME IN *IPOMOEA LACUNOSA*": Rifkin_Appendix_S15_bias_vs_PVE_RHE

Appendix S14. The relationships of bias and absolute value of bias (bias magnitude) with average total percent variance explained (PVE) and average total relative homozygous effect (RHE) for all pairs of traits.

Analyses used all QTLs. None of the correlations are significant by permutation test.

A	B

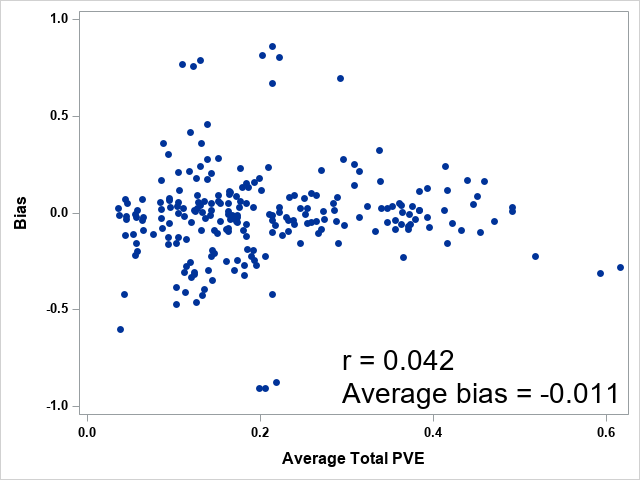

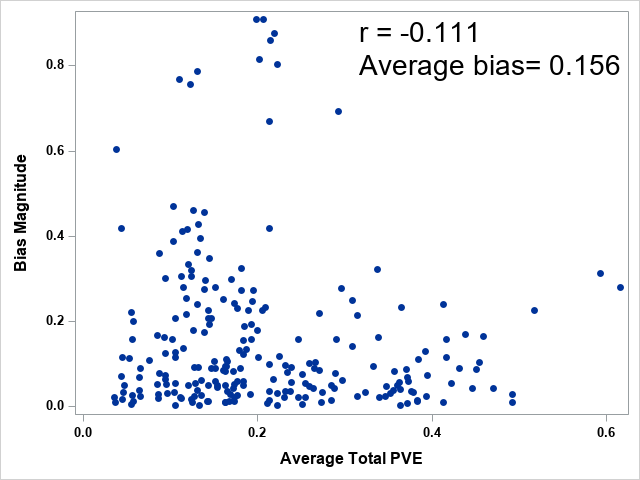

C	D

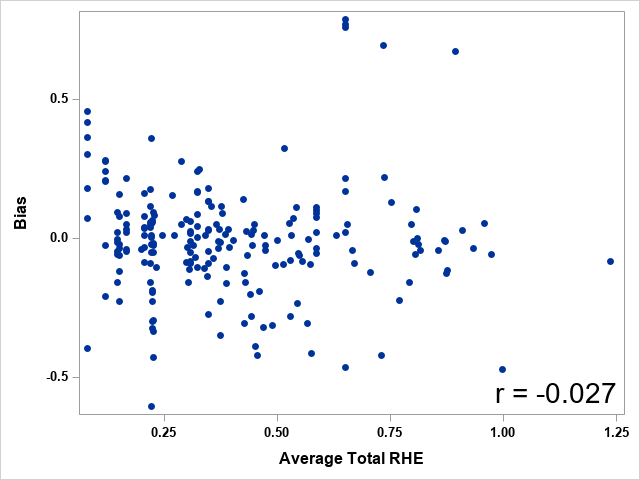

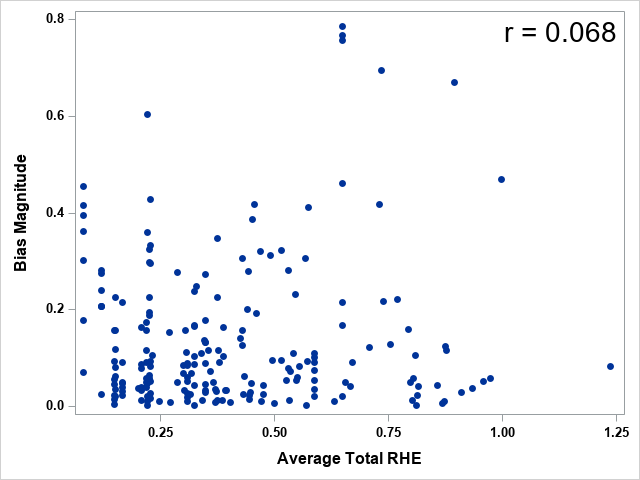
