## Supplementary material for "GENETIC ARCHITECTURE OF DIVERGENCE: THE SELFING SYNDROME IN *IPOMOEA LACUNOSA*": Rifkin_Appendix_S16_bias_vs_PVE_RHE_GWS_only

Appendix S15. The relationships of bias and absolute value of bias (bias magnitude) with average total percent variance explained (PVE) and average total relative homozygous effect (RHE) for all pairs of traits.

Analyses used only GWS QTLs. None of the correlations are significant by permutation test.

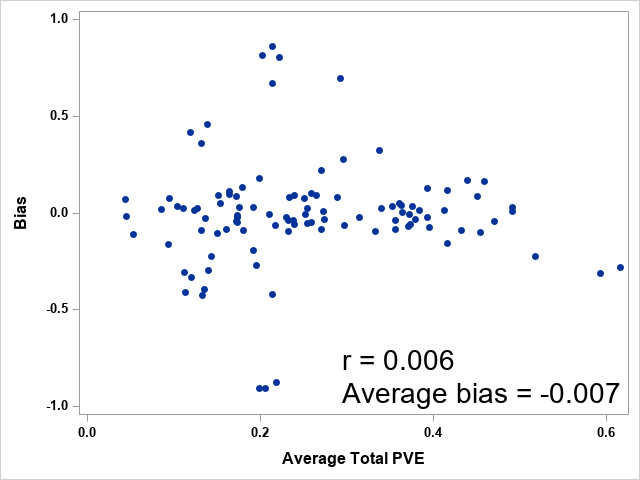

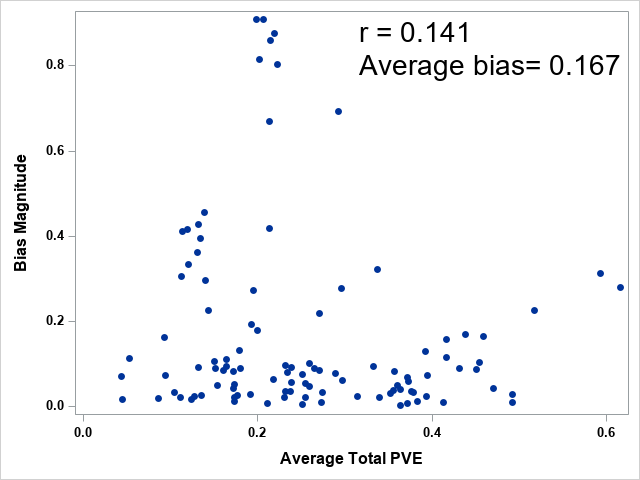

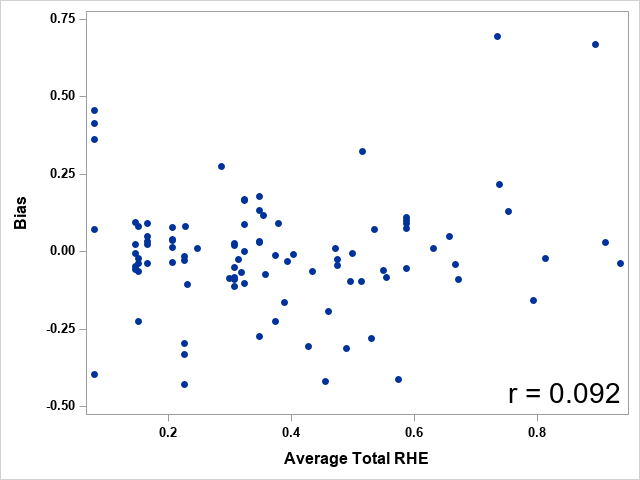

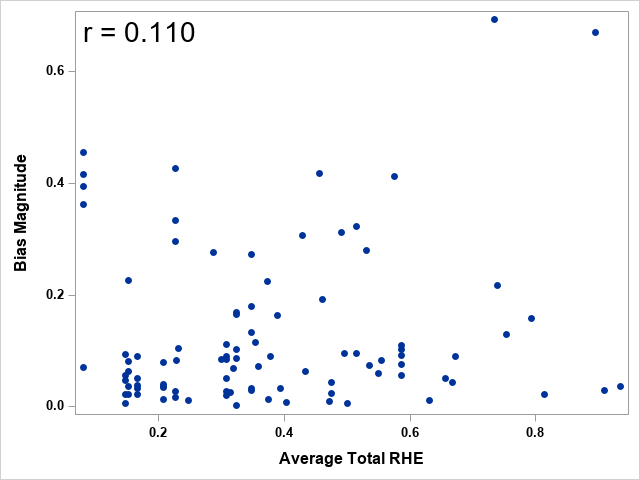
