## Supplementary material for "GENETIC ARCHITECTURE OF DIVERGENCE: THE SELFING SYNDROME IN *IPOMOEA LACUNOSA*": Rifkin_Appendix_S17_anatomical_module_correlations

### Appendix S16 Average pairwise trait correlations (and standard error) within (main diagonal) and between (off-diagonal) modules for Anatomical modules.

Within module averages calculated as the average of correlations for all possible trait pairs within the module. Between module averages calculated as average of correlations between a trait in one module and a trait in the second module. Mean within-module correlation = 0.386. Mean between-module correlation = 0.111. Difference significant by permutation test (P < 0.001). There is no standard error for the within module 5 entry because there are only two traits in this module.

|  | Flower morphology | Inflorescence | Phenology | Pollen | Nectar |
| --- | --- | --- | --- | --- | --- |
| Flower morophology | 0.440 (0.053) | 0.142 (0.041) | 0.059 (0.006) | 0.078 (0.011) | 0.314 (0.046) |
| Inflorescence |  | 0.535 (--) | 0.054 (0.019) | 0.083 (0.032) | 0.236 (0.111) |
| Phenology |  |  | 0.350 (0.032) | 0.197 (0.024) | 0.068 (0.021) |
| Pollen |  |  |  | 0.473 (0.079) | 0.041 (0.032) |
| Nectar |  |  |  |  | -- |
