## Supplementary material for "GENETIC ARCHITECTURE OF DIVERGENCE: THE SELFING SYNDROME IN *IPOMOEA LACUNOSA*": Rifkin_Appendix_S18_QTL_overlap_anatomical_modules

### Appendix S17 Average QTL overlap (and standard error) within and between anatomical modules.

### A. Only QTLs exhibiting genome-wide significance. Average overlap (and standard error) within modules = 0.377 (0.060). Average overlap between modules = 0.007 (0.004). Difference significant at P < 0.001 (permutation test) B. All QTLs. Average overlap (and standard error) within modules = 0.244 (0.031). Average overlap between modules = 0.052 (0.007). Difference significant at P < 0.001 (permutation test). NA: estimate not available because no, or only 1, data point.

A. Genome-wide significant QTLs

|  | Floral morphology | Inflorescence | Phenology | Pollen | Nectar |
| --- | --- | --- | --- | --- | --- |
| Floral morphology | 0.259 (0.085) | 0.028 (0.028) | 0.000 (0.000) | 0.000 (0.000) | 0.056 (0.035) |
| Inflorescence |  | NA | 0.000 (0.000) | 0.000 (0.000) | 0.000 (0.000) |
| Phenology |  |  | 0.494 (0.076) | 0.000 (0.000) | 0.000 (0.000) |
| Pollen |  |  |  | NA | 0.000 (NA) |
| Nectar |  |  |  |  | NA |

B. All QTLs (genome-wide and chromosome-wide significance)

|  | Floral morphology | Inflorescence | Phenology | Pollen | Nectar |
| --- | --- | --- | --- | --- | --- |
| Floral morphology | 0.305 (0.070) | 0.130 (0.028) | 0.037 (0.009) | 0.047 (0.015) | 0.284 (0.075) |
| Inflorescence |  | 0.333 (NA) | 0.031 (0.009) | 0.000 (0.000) | 0.167 (0.167) |
| Phenology |  |  | 0.208 (0.036) | 0.013 (0.009) | 0.000 (0.000) |
| Pollen |  |  |  | 0.444 (0.056) | 0.000 (NA) |
| Nectar |  |  |  |  | NA |
