## Supplementary material for "GENETIC ARCHITECTURE OF DIVERGENCE: THE SELFING SYNDROME IN *IPOMOEA LACUNOSA*": Rifkin_Appendix_S19_QTL_localization

Appendix S18 Results of randomization test for QTL co-localization for cluster modules.

QTLs were randomly assigned to positions in genome and overlaps calculated for 1,000 replicates.

P is proportion of randomized trials with average overlap ≥ observed. “All” indicates averaging over all modules. NA indicates average overlap not calculable because the are QTLs for only one trait. A. All QTLs. B. Only GWS QTLs.

**A. All QTLs**

Module	Observed

Average Overlap	P

1	0.2987	0

2	0.1778	0

3	0.3667	0

4	0.2222	0

5	0.4517	0

All	0.319	0

**B. Only GWS QTLs**

Module	Observed	Proportion of Randomized trials

Average Overlap	with Average overlap ≥ observed

1	0.2001	0

2	NA	NA

3	1.00	0.066

4	NA	NA

5	0.625	0

All	0.320	0
