## Supplementary material for "GENETIC ARCHITECTURE OF DIVERGENCE: THE SELFING SYNDROME IN *IPOMOEA LACUNOSA*": Rifkin_Appendix_S20_predicted_correlation_correlations_anatomical

Appendix S19. Relationship between average QTL overlap and average phenotypic correlations for anatomical modules.

A. All QTLs. Crorrelation significant by permutation test (P = 0.025). B. Only GWS QTLs. Correlation not significant by permutation test (P = 0.190).

A	B

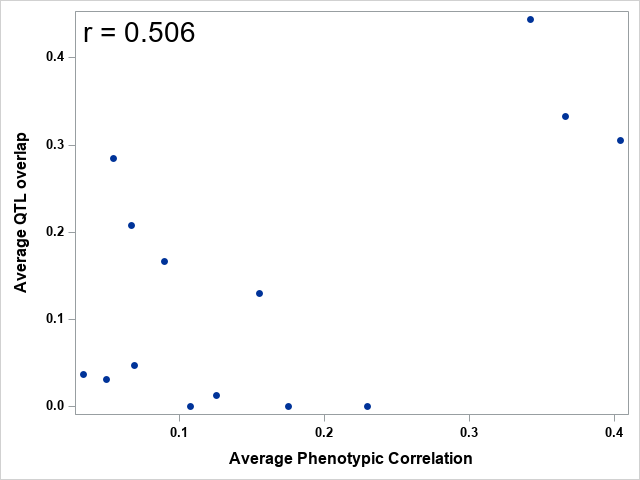

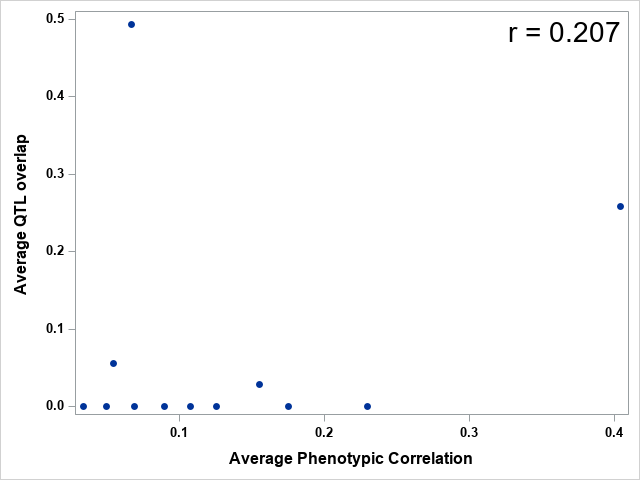
