## Supplementary material for "GENETIC ARCHITECTURE OF DIVERGENCE: THE SELFING SYNDROME IN *IPOMOEA LACUNOSA*": Rifkin_Appendix_S21_Quant_review_supp_table

Appendix S20 Jaccard overlaps of modules in selfing syndrome QTL studies

Lin and Ritland, 1997

*Mimulus guttatus* and *M. platycalyx* (Phrymaceae)

|  | Floral | Color | Inflorescence | Phenology | Nectar | Pollen | Reproductive | Vegetative |
| --- | --- | --- | --- | --- | --- | --- | --- | --- |
| Floral | 0.144 |  |  |  |  |  | 0.087 |  |
| Color |  |  |  |  |  |  |  |  |
| Inflorescence |  |  |  |  |  |  |  |  |
| Phenology |  |  |  |  |  |  |  |  |
| Nectar |  |  |  |  |  |  |  |  |
| Pollen |  |  |  |  |  |  |  |  |
| Reproductive |  |  |  |  |  |  | N/A |  |
| Vegetative |  |  |  |  |  |  |  |  |

Fishman and Willis, 2002

*Mimulus guttatus* and *M. nasutus* (Phrymaceae)

|  | Floral | Color | Inflorescence | Phenology | Nectar | Pollen | Reproductive | Vegetative |
| --- | --- | --- | --- | --- | --- | --- | --- | --- |
| Floral | 0.343 |  |  |  |  |  |  |  |
| Color |  |  |  |  |  |  |  |  |
| Inflorescence |  |  |  |  |  |  |  |  |
| Phenology |  |  |  |  |  |  |  |  |
| Nectar |  |  |  |  |  |  |  |  |
| Pollen |  |  |  |  |  |  |  |  |
| Reproductive |  |  |  |  |  |  |  |  |
| Vegetative |  |  |  |  |  |  |  |  |

Fishman et al. 2014

*Mimulus lewisii* and *M. parishii* (Phrymaceae)

|  | Floral | Color | Inflorescence | Phenology | Nectar | Pollen | Reproductive | Vegetative |
| --- | --- | --- | --- | --- | --- | --- | --- | --- |
| Floral | 0.366 |  | 0.267 | 0.215 |  |  | 0.205 |  |
| Color |  |  |  |  |  |  |  |  |
| Inflorescence |  |  | N/A | 0.286 |  |  | 0 |  |
| Phenology |  |  |  | N/A |  |  | 0 |  |
| Nectar |  |  |  |  |  |  |  |  |
| Pollen |  |  |  |  |  |  |  |  |
| Reproductive |  |  |  |  |  |  | 0.600 |  |
| Vegetative |  |  |  |  |  |  |  |  |

Bernacchi and Tanksley, 1999

*Solanum habrochaites* and *S. esculentum* (Solanaceae)

|  | Floral | Color | Inflorescence | Phenology | Nectar | Pollen | Reproductive | Vegetative |
| --- | --- | --- | --- | --- | --- | --- | --- | --- |
| Floral | 0.234 |  | 0.202 | 0.271 |  |  | 0.098 |  |
| Color |  |  |  |  |  |  |  |  |
| Inflorescence |  |  | 0 | 0 |  |  | 0.133 |  |
| Phenology |  |  |  | 0 |  |  | 0.667 |  |
| Nectar |  |  |  |  |  |  |  |  |
| Pollen |  |  |  |  |  |  |  |  |
| Reproductive |  |  |  |  |  |  | 0.333 |  |
| Vegetative |  |  |  |  |  |  |  |  |

Georgiady et al., 2002

*Solanum pimpinellifolium* (Solanaceae)

|  | Floral | Color | Inflorescence | Phenology | Nectar | Pollen | Reproductive | Vegetative |
| --- | --- | --- | --- | --- | --- | --- | --- | --- |
| Floral | 0.167 |  | 0.5 |  |  |  |  |  |
| Color |  |  |  |  |  |  |  |  |
| Inflorescence |  |  | N/A |  |  |  |  |  |
| Phenology |  |  |  |  |  |  |  |  |
| Nectar |  |  |  |  |  |  |  |  |
| Pollen |  |  |  |  |  |  |  |  |
| Reproductive |  |  |  |  |  |  |  |  |
| Vegetative |  |  |  |  |  |  |  |  |

Goodwillie et al. 2006

*Leptosiphon bicolor* and *L. jepsonii* (Polemoniaceae)

|  | Floral | Color | Inflorescence | Phenology | Nectar | Pollen | Reproductive | Vegetative |
| --- | --- | --- | --- | --- | --- | --- | --- | --- |
| Floral | 0.301 |  |  |  |  |  |  |  |
| Color |  |  |  |  |  |  |  |  |
| Inflorescence |  |  |  |  |  |  |  |  |
| Phenology |  |  |  |  |  |  |  |  |
| Nectar |  |  |  |  |  |  |  |  |
| Pollen |  |  |  |  |  |  |  |  |
| Reproductive |  |  |  |  |  |  |  |  |
| Vegetative |  |  |  |  |  |  |  |  |

Sicard et al. 2011

*Capsella rubella* and *C. grandiflora* (Brassicaceae)

|  | Floral | Color | Inflorescence | Phenology | Nectar | Pollen | Reproductive | Vegetative |
| --- | --- | --- | --- | --- | --- | --- | --- | --- |
| Floral | 0.286 |  |  |  |  |  | 0.225 | 0.208 |
| Color |  |  |  |  |  |  |  |  |
| Inflorescence |  |  |  |  |  |  |  |  |
| Phenology |  |  |  |  |  |  |  |  |
| Nectar |  |  |  |  |  |  |  |  |
| Pollen |  |  |  |  |  |  |  |  |
| Reproductive |  |  |  |  |  |  | N/A | 0.333 |
| Vegetative |  |  |  |  |  |  |  | 1 |

Slotte et al. 2012

*Capsella rubella* and *C. grandiflora* (Brassicaceae)

|  | Floral | Color | Inflorescence | Phenology | Nectar | Pollen | Reproductive | Vegetative |
| --- | --- | --- | --- | --- | --- | --- | --- | --- |
| Floral | 0.277 |  |  | 0.147 |  |  | 0.227 |  |
| Color |  |  |  |  |  |  |  |  |
| Inflorescence |  |  |  |  |  |  |  |  |
| Phenology |  |  |  | 0.333 |  |  | 0.036 |  |
| Nectar |  |  |  |  |  |  |  |  |
| Pollen |  |  |  |  |  |  |  |  |
| Reproductive |  |  |  |  |  |  | 0.242 |  |
| Vegetative |  |  |  |  |  |  |  |  |

Wozniak et al. 2019

*Capsella orientalis* and *C. grandiflora* (Brassicaceae)

|  | Floral | Color | Inflorescence | Phenology | Nectar | Pollen | Reproductive | Vegetative |
| --- | --- | --- | --- | --- | --- | --- | --- | --- |
| Floral | 0.369 |  |  |  |  |  | 0.081 | 0.101 |
| Color |  |  |  |  |  |  |  |  |
| Inflorescence |  |  |  |  |  |  |  |  |
| Phenology |  |  |  |  |  |  |  |  |
| Nectar |  |  |  |  |  |  |  |  |
| Pollen |  |  |  |  |  |  |  |  |
| Reproductive |  |  |  |  |  |  | 0.200 | 0.143 |
| Vegetative |  |  |  |  |  |  |  | N/A |

Wozniak et al. 2019

*Capsella rubella* and *C. grandiflora* (Brassicaceae)

|  | Floral | Color | Inflorescence | Phenology | Nectar | Pollen | Reproductive | Vegetative |
| --- | --- | --- | --- | --- | --- | --- | --- | --- |
| Floral | 0.222 |  |  |  |  |  | 0.194 |  |
| Color |  |  |  |  |  |  |  |  |
| Inflorescence |  |  |  |  |  |  |  |  |
| Phenology |  |  |  |  |  |  |  |  |
| Nectar |  |  |  |  |  |  |  |  |
| Pollen |  |  |  |  |  |  |  |  |
| Reproductive |  |  |  |  |  |  | N/A |  |
| Vegetative |  |  |  |  |  |  |  |  |
