## Supplementary material for "GENETIC ARCHITECTURE OF DIVERGENCE: THE SELFING SYNDROME IN *IPOMOEA LACUNOSA*": Rifkin_Appendix_S1_Measurement_methods

357 F2 individuals flowered and were phenotyped for ten floral and inflorescence traits: color (COL), number of flowers open (FPD), focal cyme length (CY), number of flowers or buds on focal cyme (FC), corolla length (CL), corolla width (CW), corolla tissue length (TL), herkogamy (HK), nectar volume (NVL), and style length (SL). Between one and 27 individuals were phenotyped daily (mean 11.3) between July 8 2014 and March 17 2015. A single open flower from a flowering individual was selected at random until at least 5 flowers had been measured (mean 3.76 flowers per individual). Floral measurements for each individual were averaged across all flowers for analysis. Unless otherwise noted, all measurements were performed using a digital caliper (Mitutoyo Digimatic CD6” CS).

Floral measurements were performed as follows:

Color was scored qualitatively as either “purple” or “white.” Colors were coded numerically as purple=1 and white=0.

The total number of flowers open on the individual flower was recorded (FPD in Table 1). The cyme bearing the selected flower was measured (cyme length, CY) as was the total number of flowers or buds on that cyme (flowers or buds per cyme, FC).

We measured corolla length (CL), corolla width (CW) and corolla tissue length (TL) from intact flowers, starting at the base of the ovary, as shown in Figure S1 below. Because *I. lacunosa* produces narrower flowers than I. cordatotriloba (Duncan and Rausher 2013a), we also included two derived measures of flower shape: CL/CW (LW) and TL/CL (TLL). Style length (SL) was measured on dissected flowers from the base of the ovary to the tip of the stigma (i.e. the entire gynoecium; the boundary between the ovary and the style is not clearly delimited).

We measured herkogamy (HK) with a five-point scale adapted from (Duncan and Rausher 2013a). The points were -1 (stigma inserted below all anthers), -0.5 (stigma below anthers, 1 anther touching stigma), 0 (2-5 anthers touching stigma), 0.5 (stigma above anthers, 1 anther touching stigma), and 1 (stigma exserted above all anthers).

To measure nectar volume (NVL), we applied a capillary tube (Drummond, 0.4mm diameter) to the nectary of a dissected flower and measured the height of nectar drawn up with a digital caliper. Height was converted to volume using the formula for the volume of a cylinder (V=πr2h).

We used two approaches to quantify pollen number (P1, P2) and size (PS) from a subset of F2 individuals (N = 110 number lactophenol and ImageJ (P2), N=69 number Coulter Counter (P1), N=68 size) and the parent and grandparents. For P2, we collected anthers from unopened buds the day before anthesis and stained them with lactophenol, then counted them using ImageJ software (Schindelin et al. 2012) following the approach of (Costa and Yang 2009). Pollen diameter (Pollen Size (PS) in Table 1) and an additional measure of pollen number (Pollen Count 1 (P1) in Table 1) were obtained using a Coulter Z2 counter. Because pollen quantification sampled unopened flowers destructively, we took pollen data only from a subset of plants that produced many flowers, which likely biases our results in assessing pollen traits, although in an unknown direction.

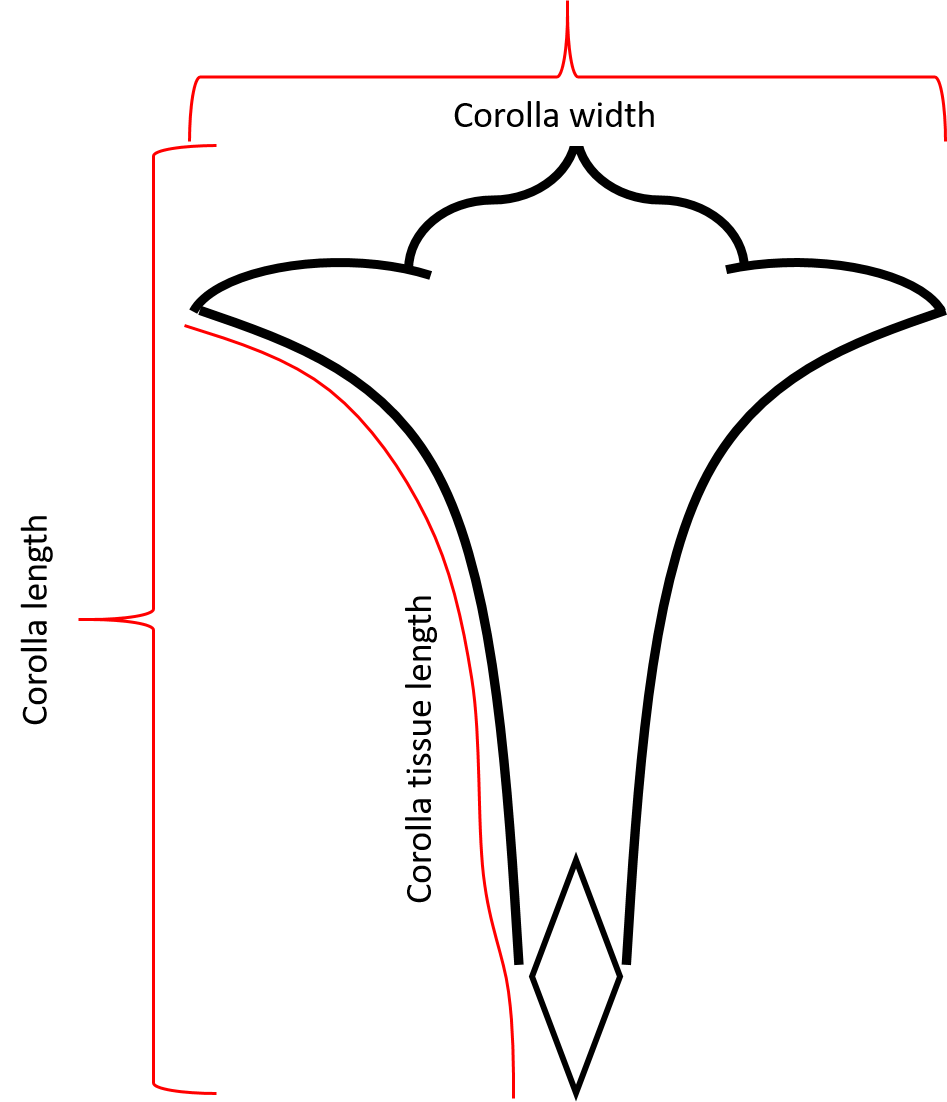

Figure S1

Floral morphological measurements of corolla length, corolla width and corolla tissue length.
