## Supplementary material for "GENETIC ARCHITECTURE OF DIVERGENCE: THE SELFING SYNDROME IN *IPOMOEA LACUNOSA*": Rifkin_Appendix_S2_ddRADseq_library_construction_and_sequencing

We generated markers from our mapping population using double-digest restriction-site associated DNA sequencing (ddRADseq (Peterson et al. 2012)). DNA was isolated from flash-frozen young leaf tissue homogenized with a GenoGrinder (SPEX Sample Prep) using ThermoFisher GeneJet Plant DNA kits. We digested samples with EcoR1 and MSP1 (New England Biolabs) and cleaned samples with AMPure XP beads (Agencourt). DNA was quantified for optimizing ligation efficiency and for library balancing using a Qubit dsDNA BR assay kit (Thermo Fisher). 96 eight-base in-line forward barcodes (Parchman et al. 2011) were ligated to 100ng DNA per sample with T4 DNA ligase (New England Biolabs). We pooled our samples into five plates for size selection at the Duke Sequencing and Genomic Technologies core resource. Samples were amplified after pooling. We initially sequenced two lanes on the Illumina HiSeq platform, then compensated for stochastic differences in amplification by resequencing the 192 lowest-coverage individuals in two successive rounds of one and two lanes respectively. Finally, 33 samples were included in a later 4-lane sequencing project for more coverage.

In preliminary analyses, one half-plate from the library construction consistently showed large numbers of double crossovers. We determined that this was suggestive of contamination and therefore omitted those samples from subsequent analyses. A subset of samples from that half-plate and from other individuals that had had low coverage in the initial sequencing run were added to a subsequent sequencing run. We ultimately included 396 F2 individuals in our linkage map and subsequent QTL analysis.
