## Supplementary material for "GENETIC ARCHITECTURE OF DIVERGENCE: THE SELFING SYNDROME IN *IPOMOEA LACUNOSA*": Rifkin_Appendix_S3_Linkage map construction

A linkage map was constructed using Lep-Map3 (Rastas et al., 2013) from reads aligned to the *I. lacunosa* draft assembly. ddRADseq sequencing reads were aligned to the draft genome using NextGenMap (Sedlazeck et al., 2013) and cleaned, sorted, and assigned read groups using Picard Tools 2.18.21-SNAPSHOT 568 (https://broadinstitute.github.io/picard/). Genotype probabilities were called using the Lep-Map3 pileupParser2 module, resulting in an initial 393,900 loci, and related to parental genotypes using the ParentCall2 module, resulting in 15,426 loci. Markers were separated into linkage groups with an initial LOD cutoff of 53, resulting in 11 major linkage groups (>230 markers). The largest linkage group (2450 markers) was then split with a higher LOD cutoff (95), resulting in 15 major linkage groups of 120-762 markers, consistent with the karyotype of *I. lacunosa* (Nakajima, 1963), and 4 minor linkage groups of 10-59 markers. Markers were ordered using the OrderMarkers2 module of Lep-Map3 and phased in selfing mode.

We then used Lep-Anchor (Rastas, 2020) to relate our linkage map to our draft assembly and generate marker orders that incorporated the physical and linkage estimates. This resulted in 17 linkage groups of 50-694 markers. We constructed Marey maps (Chakravarti, 1991) in R to examine collinearity between the linkage map and the draft assembly. Marker positions were generally highly consistent between the linkage map and draft assembly. We manually removed markers that caused significant map expansion, which likely reflect genotyping error. Recombination plots and Marey maps revealed that two of the draft assembly scaffolds had been split, so we manually rejoined them in the .agp files before producing the final maps. We generated the final maps by positioning markers according to Lep-Anchor’s integration of the draft assembly and the linkage data. Scripts are available on Github [XXX]

Chakravarti, A. 1991. A graphical representation of genetic and physical maps: the Marey map. *Genomics* 11: 219–222.

Nakajima, G. 1963. Karyotype of Genus Ipomoea. *Cytologia* 28: 351–359.

Rastas, P. 2020. Lep-Anchor: automated construction of linkage map anchored haploid genomes. *Bioinformatics (Oxford, England)* 36: 2359–2364.

Rastas, P., L. Paulin, I. Hanski, R. Lehtonen, P. Auvinen, and M. Brudno. 2013. Lep-MAP: Fast and accurate linkage map construction for large SNP datasets. *Bioinformatics* 29: 3128–3134.

Sedlazeck, F. J., P. Rescheneder, and A. Von Haeseler. 2013. NextGenMap: Fast and accurate read mapping in highly polymorphic genomes. *Bioinformatics* 29: 2790–2791.
