## Supplementary material for "GENETIC ARCHITECTURE OF DIVERGENCE: THE SELFING SYNDROME IN *IPOMOEA LACUNOSA*": Rifkin_Appendix_S4_QTL_mapping

### Appendix S5. QTL mapping.

We generated our genotype data for QTL mapping using Lep-Map3’s map2genotypes function and custom scripts to identify grandparent genotypes from the output of parentcall2. Single-QTL analyses were performed in QTL2 (Broman et al., 2019). For QTL mapping, we first examined the distributions of traits. Normally-distributed and binary traits were not transformed. Skewed traits were transformed or split into two-part models, as recommended in the R/QTL forum (https://groups.google.com/forum/#!searchin/rqtl-disc/2-part%7Csort:date/rqtl-disc/7dHDVAedcmc/1FUFZh6qCAAJ).

The following traits were approximately normally distributed: corolla length (CL), corolla width (CW), corolla tissue length (TL), date of flower measurement (FD), height in mm on day 21 (H), length of first three internodes on day 21 (INT1, INT2, INT3), both measures of corolla shape (LW, TLL), number of leaves on day 21 (NL), how many flowers were measured (NF), cyme length (CY), both measures of pollen count (P1, P2), pollen size (PS), and style length (SL). The following traits were binary: color (C), whether a plant ever produced flowers (EF), and sterile morphology (ST). Nectar volume (NVL) was log-transformed (1 was added to all values first because it was zero-inflated), number of flowers per day and number of flowers on cyme were both converted to two-part models (a binary model for one or more than one, and a continuous model for more than one: FC_split_continuous and FC_split_binary from FC, and FPD_binary from FPD. No QTLs were identified for FPD_continuous), and we performed a negative reciprocal transformation on dates of the first three leaves opening. We therefore identified QTLs for a total of 27 phenotypes: the 26 listed in Table 1, plus additional phenotypes for the two that were split, minus herkogamy (Appendix S4). Scripts are available on Github [https://github.com/joannarifkin/Ipomoea_QTL].

We used diagnostic features in QTL2 to identify individuals with high numbers of crossovers, suggestive of genotyping errors, and removed 10. We also used diagnostics from ASMAP (Taylor and Butler, 2017) to identify distorted markers, but did not remove them. Our final QTL mapping dataset included 386 phenotyped and genotyped individuals. To identify QTLs, we used the scan1 function in QTL2 with either the default or binary model. We determined significance cutoffs using permutation analysis (1000 replicates) at both genome-wide and chromosome-wide levels. Confidence intervals were estimated as 1.5-LOD intervals. We perform all downstream analyses separately for both “all” and “genome-only” significant QTLs. Additive and dominance effects were estimated using the scan1coef function in QTL2. QTLs were considered dominant if the dominance deviation was >0.9x the additive effect, additive if the dominance deviation was <0.1x the additive effect, and partly dominant in between. QTLs were categorized as over- or under-dominant if the heterozygote value was outside of the range of the homozygote values.

Broman, K. W., D. M. Gatti, P. Simecek, N. A. Furlotte, P. Prins, Ś. Sen, B. S. Yandell, and G. A. Churchill. 2019. R/qtl2: Software for mapping quantitative trait loci with high-dimensional data and multiparent populations. *Genetics* 211: 495–502.

Taylor, J., and D. Butler. 2017. R package ASMap: efficient genetic linkage map construction and diagnosis. *Journal of Statistical Software* 79.
