## Supplementary material for "GENETIC ARCHITECTURE OF DIVERGENCE: THE SELFING SYNDROME IN *IPOMOEA LACUNOSA*": Rifkin_Appendix_S6_Genome_annotation

A de novo annotation of *Ipomoea lacunosa* genome was generated by combining evidence alignment and de-novo prediction. Repeat sequences were identified by RepeatMasker v4.1.0 and Repeat Modeler v2.0.1 (Smit and Hubley, 2008; Smit et al., 2013). For prediction, we used a transcript generated by first aligning RNA-seq sequences to the *I. lacunosa* genome (Rifkin et al., 2019) using by STAR v2.7.1a (Dobin et al., 2013), then merging the alignments into one sequence by StringTie v2.1.1 (Pertea et al., 2015). This evidence alignment prediction and *ab Initio* prediction by GeneMark-ES and Augustus v3.3.4 was achieved by BRAKER2 (Hoff et al., 2019; Brůna et al., 2020). This annotation was then used to train *ab Initio* prediction software SNAP (Korf, 2004). The output of SNAP was introduced into the annotation by MAKER v2.31.10 (Cantarel et al., 2008). This step to train and predict was iterated for 3 times, and the final structure annotation was created by MAKER. After each iteration, the AED score was checked, and annotation statistics was measured by agat v0.4.0 (Dainat, n.d.). The output gff file was also trimmed to remove duplications and broken genes by agat v0.4.0. The best structure prediction was selected based on AED score and agat report. This prediction was searched against for function annotation with eggNOG-mapper (Huerta-Cepas et al., 2017, 2019).

Brůna, T., K. J. Hoff, A. Lomsadze, M. Stanke, and M. Borodovsky. 2020. BRAKER2: Automatic Eukaryotic Genome Annotation with GeneMark-EP+ and AUGUSTUS Supported by a Protein Database. 1–21.

Cantarel, B. L., I. Korf, S. M. C. Robb, G. Parra, E. Ross, B. Moore, C. Holt, et al. 2008. MAKER: An easy-to-use annotation pipeline designed for emerging model organism genomes. *Genome Research* 18: 188–196.

Dainat, J. AGAT: Another Gff Analysis Toolkit to handle annotations in any GTF/GFF format. (Version v0.4.0). Zenodo. Website https://www.doi.org/10.5281/zenodo.3552717.

Dobin, A., C. A. Davis, F. Schlesinger, J. Drenkow, C. Zaleski, S. Jha, P. Batut, et al. 2013. {STAR}: ultrafast universal {RNA-seq} aligner. *Bioinformatics* 29: 15–21.

Hoff, K. J., A. Lomsadze, M. Borodovsky, and M. Stanke. 2019. Whole-genome annotation with BRAKER. *In* M. Kollmar [ed.], Gene Prediction Methods and Protocols Methods in Molecular Biology, 65–96. Humana Press, New York.

Huerta-Cepas, J., K. Forslund, L. P. Coelho, D. Szklarczyk, L. J. Jensen, C. Von Mering, and P. Bork. 2017. Fast genome-wide functional annotation through orthology assignment by eggNOG-mapper. *Molecular Biology and Evolution* 34: 2115–2122.

Huerta-Cepas, J., D. Szklarczyk, D. Heller, A. Hernández-Plaza, S. K. Forslund, H. Cook, D. R. Mende, et al. 2019. EggNOG 5.0: A hierarchical, functionally and phylogenetically annotated orthology resource based on 5090 organisms and 2502 viruses. *Nucleic Acids Research* 47: D309–D314.

Korf, I. 2004. Gene finding in novel genomes. *BMC Bioinformatics* 5: 1–9.

Pertea, M., G. M. Pertea, C. M. Antonescu, T.-C. Chang, J. T. Mendell, and S. L. Salzberg. 2015. StringTie enables improved reconstruction of a transcriptome from RNA-seq reads. *Nature Biotechnology* 33: 290–295.

Rifkin, J. L., A. S. Castillo, I. T. Liao, and M. D. Rausher. 2019. Gene flow, divergent selection and resistance to introgression in two species of morning glories (*Ipomoea*). *Molecular Ecology* 28: 1709–1729.

Smit, A. F. A., and R. Hubley. 2008. RepeatModeler Open-1.0. Website http://www.repeatmasker.org.

Smit, A. F. A., R. Hubley, and P. Green. 2013. RepeatMasker Open-4.0. Website http://www.repeatmasker.org.
