## Supplementary material for "GENETIC ARCHITECTURE OF DIVERGENCE: THE SELFING SYNDROME IN *IPOMOEA LACUNOSA*": Rifkin_Appendix_S7_Predicting_Genetic_Correlations_from_QTL_properties

### Appendix S6. Predicting Genetic Correlations from QTL properties.

We assessed the degree to which QTL effects can predict pairwise genetic correlations between traits, using a modification of the approach described by Gardner and Latta (2007). Mean phenotypic values for each genotype at a QTL locus were obtained using QTL2. For each trait *i*, we calculated the genetic variance associated with each QTL *j* as

${Var}_{{QTL}_{ij}}= p_{11}\left( a_{ij}-\mu_{ij} \right)^{2}+p_{12}\left( d_{ij}-\mu_{ij} \right)^{2}+ p_{22}\left( b_{ij}-\mu_{ij} \right)^{2}$

where *p*_11_, *p*_12_, and *p*_22_ are the frequencies of individuals homozygous for the *I. cordatotriloba* allele, heterozygous, and homozygous for the *I. lacunosa* allele respectively. $a_{ij}$ is the average trait value of homozygotes for the *I. cordatotriloba* allele, $b_{ij}$ is the analogous value for homozygotes for the *I. lacunosa* allele, and $d_{ij}$ is the average trait value for heterozygotes. Finally, $\mu_{ij}$ is the average value of the trait, i.e.

$$\mu_{ij}=p_{11}a_{ij}+p_{12}d_{ij}+ p_{22} b_{ij}$$

The total genetic variance contributed to character *i* by all QTLs is then

$${Var}_{G_{i}}= \sum_{j} {Var}_{{QTL}_{ij}}$$

Similarly, for a pair of traits *i* and *k*, the genetic covariance associated with a particular co-localizing QTL *j* is

$${Cov}_{{QTL}_{ikj}}=p_{11}\left( a_{ij}-\mu_{ij} \right)\left( a_{kj}-\mu_{kj} \right)+p_{12}\left( d_{ij}-\mu_{ij} \right)\left( d_{kj}-\mu_{kj} \right)+ p_{22}\left( b_{ij}-\mu_{ij} \right)\left( b_{kj}-\mu_{kj} \right)$$

and the total covariance for the pair of traits is

${Cov}_{G_{ik}}= \sum_{j} {Cov}_{{QTL}_{ikj}}$

Finally, the predicted genetic correlation between two traits is

$r_{G_{ij}}= \frac{{Cov}_{G_{ij}}}{\sqrt{{Var}_{G_{i}} {Var}_{G_{j}}}}$

For non-co-localizing QTLs, we used the values of *p*_ij_ corresponding to the marker with the highest LOD score. For co-localizing QTLs, we assumed that the locations of causal variant is identical, which means that the QTLs for the two traits have the same value of *p*_ij_ . Because in general the marker with the highest LOD score is not identical for the two QTLs, as an approximation to the joint *p*_ij_ of two traits, we identified the midpoint of the overlap interval and used the allele frequencies from the marker closest to the midpoint.

To assess how well the predicted genetic correlations matched the actual genetic architecture of divergence, we compared the predicted correlations with the observed phenotypic correlations by calculating the correlation coefficient between the predicted and phenotypic correlations across trait pairs. Ideally, we would compare them to actual genetic correlations, but such correlations are unavailable. However, phenotypic and genetic correlations are typically highly correlated, such that phenotypic correlations can be used as a proxy for genetic correlations (Cheverud, 1988; Roff, 1995; Waitt and Levin, 1998). We also calculated bias, which is the difference between the phenotypic and predicted genetic correlations (Gardner and Latta, 2007)(Gardner and Latta, 2007). Finally, as an indicator of how completely we identified QTLs for a particular trait, we calculated total percent variance explained (PVE) by summing the PVE values for individual QTLs. As a similar indicator, we also calculated total relative homozygous effect (RHE) for traits for which we had information on the parental values. RHE for a QTL is the difference between the mean phenotypes of the two homozygotes divided by the parental difference. RHE values for individual QTLs were summed to obtain total RHE for a trait.

To determine if bias is related to completeness of QTL sampling, we calculated the correlations between bias and total PVE and between bias and total RHE. The significance of all correlations was determined using permutation analysis. For each of 1,000 permutations, we reassigned QTLs to traits randomly, calculated the variables (e.g. predicted genetic correlation, bias), and calculated the correlations. The probability of obtaining the observed correlation by chance was taken to be the proportion of permutations in which the correlations were greater than or equal to the observed correlation.

Cheverud, J. M. 1988. A comparison of genetic and phenotypic correlations. *Evolution* 42: 958–968.

Gardner, K. M., and R. G. Latta. 2007. Shared quantitative trait loci underlying the genetic correlation between continuous traits. *Molecular Ecology* 16: 4195–4209.

Roff, D. A. 1995. The estimation of genetic correlations from phenotypic correlations: A test of cheverud’s conjecture. *Heredity* 74: 481–490.

Waitt, D. E., and D. A. Levin. 1998. Genetic and phenotypic correlations in plants: A botanical test of Cheverud’s conjecture. *Heredity* 80: 310–319.
